# Depletion of microbiome-derived molecules in the host using *Clostridium* genetics

**DOI:** 10.1101/401489

**Authors:** Chun-Jun Guo, Breanna M. Allen, Kamir J. Hiam, Dylan Dodd, Will van Treuren, Steven Higginbottom, Curt R. Fischer, Justin L. Sonnenburg, Matthew H. Spitzer, Michael A. Fischbach

## Abstract

The gut microbiota produce hundreds of molecules that are present at high concentrations in circulation and whose levels vary widely among humans. In most cases, molecule production has not been linked to specific bacterial strains or metabolic pathways, and unraveling the contribution of each molecule to host biology remains difficult. A general system to ‘toggle’ molecules in this pool on/off in the host would enable interrogation of the mechanisms by which they modulate host biology and disease processes. Such a system has been elusive due to limitations in the genetic manipulability of *Clostridium* and its relatives, the source of many molecules in this pool. Here, we describe a method for reliably constructing clean deletions in a model commensal *Clostridium, C. sporogenes* (*Cs*), including multiply mutated strains. We demonstrate the utility of this method by using it to ‘toggle’ off the production of ten *Cs*-derived molecules that accumulate in host tissues. By comparing mice colonized by wild-type *Cs* versus a mutant deficient in the production of branched short-chain fatty acids, we discover a previously unknown IgA-modulatory activity of these abundant microbiome-derived molecules. Our method opens the door to interrogating and sculpting a highly concentrated pool of chemicals from the microbiome.

## MAIN TEXT

Gut bacteria produce hundreds of diffusible molecules that are notable for four reasons: 1) Most have no host source, so their levels are determined predominantly or exclusively by the microbiome. 2) Many get into the bloodstream, so they can access peripheral tissues. 3) They often reach concentrations that approach or exceed what a typical drug reaches, and the concentration range can be large – more than an order of magnitude in many cases – so they have the potential to underlie biological differences among humans. 4) Several of these molecules are known to be ligands for key host receptors; additional compounds from this category are candidate ligands for, e.g., GPCRs and nuclear hormone receptors that play an important role in the host immune and metabolic systems (*1*). Thus, the gut microbiome is a prolific endocrine organ, but its output is not well understood.

The biological activities of most of these molecules remain unknown. One reason is that there has not been a general method for ‘toggling’ one or more of them on/off in the host, akin to a gene knockout experiment in a model organism. Such a method would open the door to interrogating – and ultimately controlling – one of the most concrete contributions gut bacteria make to host biology.

Previous efforts that have sought to study an individual microbiome-derived molecule in the setting of host colonization have used two main strategies: 1) Administering a compound by injection or gavage, which can offer insights into mechanism of action but suffers from the lack of a clean background (i.e., existing physiologic levels of the molecule of interest) and the possible effects of differences in route and timing of administration relative to the native context of gut bacterial production. 2) Adding or removing a bacterial species that produces the molecule, which has the advantage of a more native-like context but makes it difficult to distinguish between molecule-induced phenotypes and other biological activities of the organism (*2–4*).

The most precise format for interrogating a microbiome-derived molecule is to compare two organisms that differ only in its production. Such an experiment has two threshold requirements: 1) knowledge of the metabolic genes for the molecule of interest, and 2) the ability to perform genetics in a robustly producing strain. This approach has been successful in *Bacteroides* (*5, 6*), *E. coli* (*7*), and *Lactobacillus* (*8*), but a key technical barrier limiting its generalization is that many of the known high-abundance gut-derived molecules are produced by *Clostridium* and its relatives, which have been difficult to manipulate genetically. We recently reported the use of a group II intron (*9*) to mutate a single pathway in *Clostridium sporogenes* (*10*), but this insertional mutagenesis system performs unpredictably and cannot be used to make strains that carry multiple mutations. Here, we address these challenges by developing a new CRISPR-Cas9-based system for constructing single and multiple mutants in a model gut-resident *Clostridium* species with high efficiency.

### Selection of *C. sporogenes* as a model gut *Clostridium*

We chose *Clostridium sporogenes* ATCC 15579 (*Cs*) as a model gut commensal from the anaerobic Firmicutes for three reasons: This strain has long been known as a robust producer of high-abundance small molecules (*11–13*), it stably colonizes germ-free mice (*3*), and it is a commensal or mutualist (i.e., neither a pathogen nor a pathobiont). We began by performing metabolic profiling experiments to determine systematically the set of high-abundance small molecules produced by this strain *in vitro*. As shown in **Figures 1** and **3**, *Cs* produces ten molecules that are highly abundant and either primarily or exclusively derived from the gut microbiota: tryptamine, indole propionate and other aryl propionates, isobutyrate, 2-methylbutyrate, isovalerate, isocaproate, propionate, and butyrate, confirming previous studies (*10–13*); and trimethylamine and 5-aminovalerate, whose production by *Cs* has not previously been reported.

**Figure 1.**
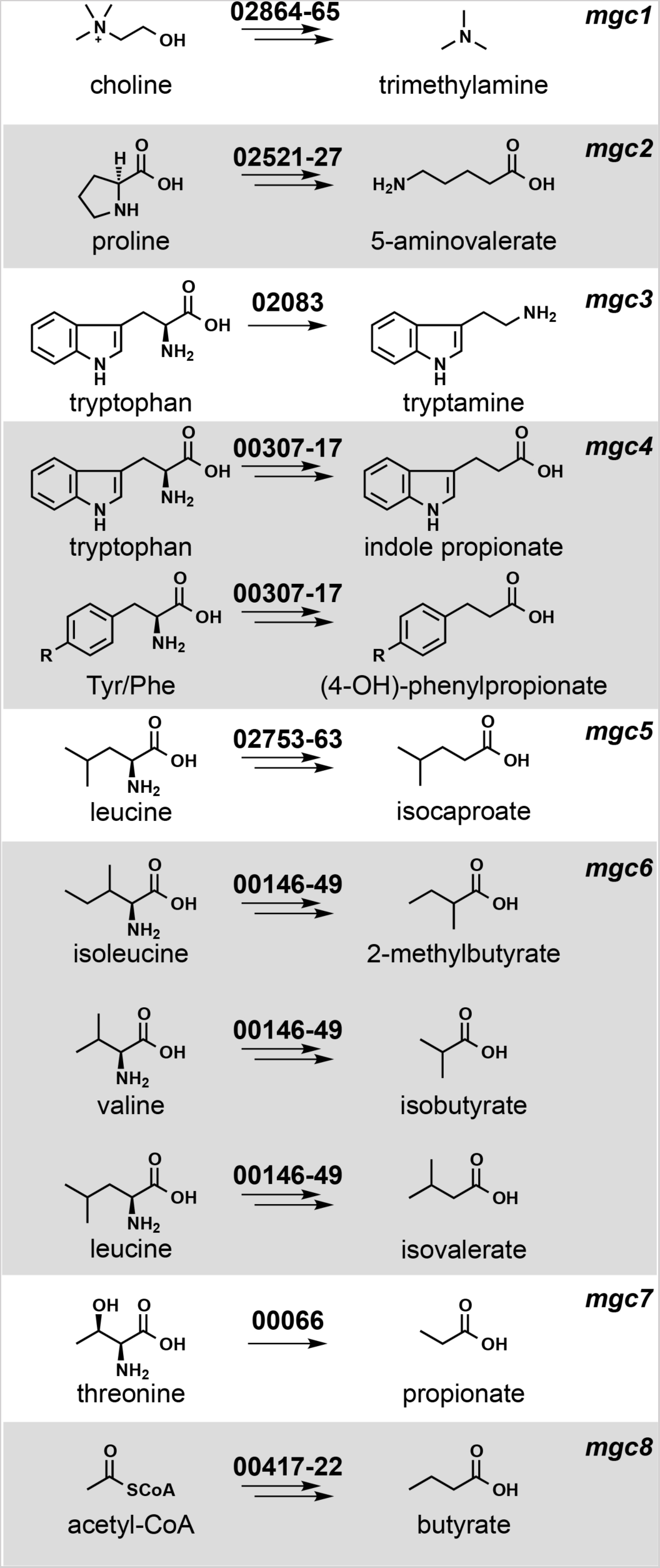
Metabolic pathways from *Clostridium sporogenes* ATCC 15579 (*Cs*) examined in this study. Each pathway generates a microbiome-derived metabolite present at high abundance in the host.The prefix “*mgc”* stands for “metabolic gene cluster”. Genes that comprise each pathway are shown above the corresponding arrows; the numbers indicate a locus tag suffix for *Cs*, where the prefix is “CLOSPO_”.

### Prediction and computational analysis of metabolic pathways

Next, we sought to predict the genes responsible for producing each molecule. Metabolic genes for trimethylamine (*14*), tryptamine (*13*), and indole propionate (*10*) have been demonstrated using genetics in *Cs* or another gut bacterium; pathways for the remaining seven molecules had not been validated genetically in the gut microbiota. We made predictions for each one based on three sources of evidence (**Figure 1**): pathways validated in non-microbiome organisms (e.g., a pathway for 5-aminovalerate production from the terrestrial isolate *Clostridium sticklandii* (*15*)); biochemical studies that implicate an enzyme superfamily, which enabled us to search for orthologs in *Cs* (e.g., 2-hydroxyisocaproate dehydratases (*16*), which led us to a predicted cluster for isocaproate); and a metagenomic analysis of butyrate gene clusters (*17*), which yielded a predicted gene cluster for butyrate production in *Cs*.

We then set out to determine whether the metabolic genes we predicted are widely distributed in the human gut microbiome and actively transcribed under conditions of colonization, reasoning that both criteria would impact the generality of our studies. We used MetaQuery (*18*) to measure the abundance of the key genes in each pathway (colored red in **Figure S1**) in >2000 publicly available human gut metagenomes. Every gene except *tdcA* was present in >95% of the stool metagenomes, indicating that the predicted pathways are cosmopolitan (minimum abundance = 1 copy/1,000 cells) (**Figure S2**). Since the mere presence of a gene in a metagenome does not imply that it is transcribed, we determined the transcript abundance of the key metabolic genes (including close homologs from non-*Cs* genomes) by recruiting reads from nine publicly available RNA-sequencing data sets derived from stool samples of healthy subjects (*19*). This analysis revealed that multiple homologs of each pathway are highly transcribed under the condition of host colonization (**Figure S2**). For example, *porA* and its homologs – ALIPUT_00387 in *Alistipes putredinis* DSM 17216, BACSTE_01839 from *Bacteroides stercoris* ATCC 43183, and BVU_2313 from *Bacteroides vulgatus* ATCC 8482 – are transcribed robustly in at least one sample. Taken together, these data suggest that the predicted pathways are widely distributed and actively transcribed under conditions of host colonization.

### Development of a new CRISPR-Cas9-based genetic system for *Cs*

Next, we tested our metabolic pathway predictions by constructing mutants in each of them, along with the known pathways in *Cs* for tryptamine (*13*), indole propionate (*10*), and trimethylamine (*14*) (**Figure 1**). The genetic system we used previously, which is based on a group II intron (*20*), had two important limitations: 1) Since the intron’s targeting mechanism is not well understood, generating one insertional mutant typically requires testing several targeting sequences; moreover, this process regularly fails to yield a mutant. 2) After multiple attempts, we were not able to recycle the antibiotic resistance marker using Clostron in order to create strains with multiple mutations; our experience is consistent with a previous report in the literature (*9*).

Reasoning that a dependable, markerless, recyclable genetic system for *Clostridium* would open the door to more systematic studies of microbiome metabolism, we developed a CRISPR/Cas9-based genome editing system for *Cs*. *Clostridium* species have been notoriously difficult to modify genetically, due in part to inefficient homologous recombination (*21*). We postulated that a Cas9-induced double-strand break would help select for a rare homologous recombination event. To this end, we constructed a single vector that includes all the essential components of a bacterial CRISPR-Cas9 system: the Cas9 gene, a guide RNA (gRNA), and a 1.5-2.5 kb repair template and transferred it by conjugation into *Cs.* However, we did not obtain viable colonies, even after multiple rounds of vector design modifications (see **Supplementary Text** for more detail).

Having observed that the conjugation efficiency for *Cs* is greatly diminished for plasmids >10 kb, we redesigned the system by splitting its components into two separate vectors: one that contains the gRNA and repair template, and another that harbors Cas9 under the control of a ferredoxin promoter (**Figures 2** and **S3**). When we introduced these plasmids sequentially, we failed to get any viable colonies after introducing the second vector that harbors Cas9 coding sequence. Reasoning that the efficiency of the second conjugation step could be the source of failure, we lengthened the donor/acceptor co-cultivation step of the second conjugal transfer from 24 to 72 h; this optimized protocol yielded reproducible, high-efficiency mutations at multiple test loci (see **Supplementary Text** for a more detailed description of the method’s development).

**Figure 2.**
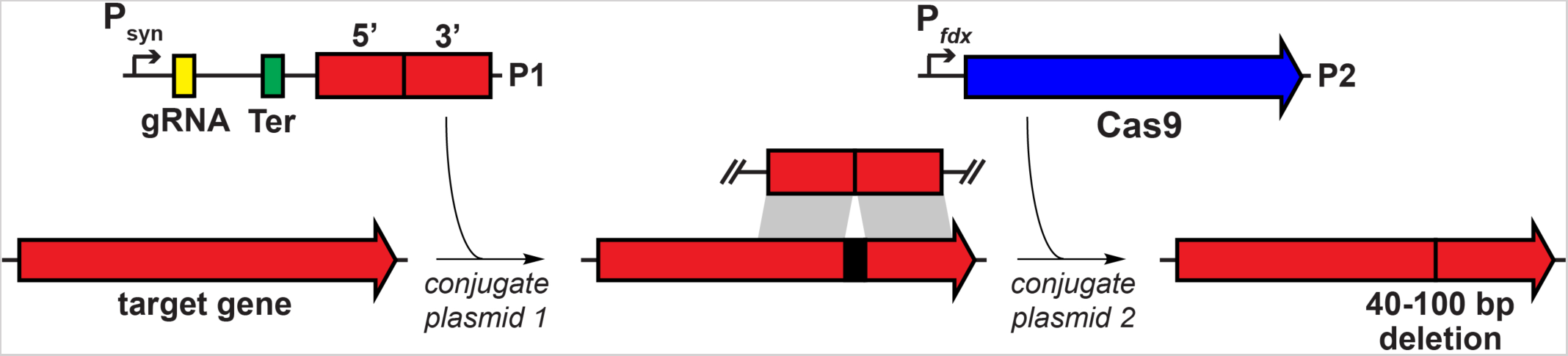
Development of a CRISPR-Cas9 based genetic system for *Cs*. Schematic view of the genetic system. In the first step, plasmid P1 is introduced by conjugation into *Cs.* P1 contains a guide RNA (gRNA) expressed under the control of Psyn, a synthetic promoter generated using PePPER (*38*); Ter, a terminator sequence from the *Cs* 16s rRNA gene; and a ∼1.5-2.0 kb repair template. In the second step, plasmid P2 is introduced by conjugation. P2 consists of the Cas9 gene from *Streptococcus pyogenes* expressed under the control of P*fdx*, the promoter from the *Cs* ferredoxin gene. Key steps in the development of the method were to introduce the genome editing components (Cas9, gRNA, and repair template) sequentially on two plasmids, and to lengthen the donor/acceptor co-cultivation step of the second conjugal transfer from 24 to 72 h.

### Constructing mutants to validate *Cs* pathways *in vitro*

We used the Cas9-based genetic system to construct deletion mutants in each of the 10 pathways. In each case, our repair template effected the removal of an 80-150 bp portion of the targeted gene in the *Cs* chromosome (**Figures 2B** and **S4**); for simplicity, the resulting mutants (e.g., *ΔporA*(330-409)) are referred to simply as *ΔporA* (see **Table S4** for a list of deleted regions). We cultured each strain in vitro and analyzed culture extracts by LC-MS or GC-MS (depending on the analyte), yielding the following conclusions: (*i*) The production of each of the ten molecules is blocked by the corresponding pathway mutant (**Figures 3** and **S5**), validating our prediction set; an important exception is detailed below in (*iii*).(*ii*) Deleting one pathway does not appreciably alter the production of other molecules, indicating these pathways function independently *in vitro*. (*iii*) Our predicted genetic locus for the branched short-chain fatty acids (branched SCFAs) isobutyrate, 2-methylbutyrate, and isovalerate proved incorrect. In other organisms (e.g., *Streptomyces avermitilis* (*22*)), the branched-chain *α*-ketoacid dehydrogenase complex (BCKDH) converts branched-chain amino acids (BCAAs) into branched SCFAs. To our surprise, deletion of the BCKDH gene CLOSPO_03305 did not affect branched SCFA production. To our surprise, the *ΔporA* mutant – a gene we previously showed to be involved in the oxidative catabolism of aromatic amino acids (*10*) – proved to be deficient in the production of all three branched SCFAs (**Figure 3**). PorA is a member of the pyruvate:ferredoxin oxidoreductase (PFOR) superfamily; like the BCKDH, PFOR enzymes are thiamine-PP dependent, but they harbor an array of iron-sulfur clusters for electron transfer in place of lipoate and flavin, and reduce ferredoxin or flavodoxin instead of NAD^+^ (*23*). Although PFOR superfamily members are known to utilize pyruvate in anaerobic bacteria (generating acetate), their utilization of branched-chain *α*-keto acids derived from BCAAs constitutes a non-canonical pathway for branched SCFA production, having only been observed in Archaea (*24, 25*). We found multiple *porA* homologs in the genomes of other human gut isolates that are highly transcribed in human stool metatranscriptomes, suggesting that this pathway could be a substantial contributor to the host branched SCFA pool (**Figures 1, 3,** and **S2**).

**Figure 3.**
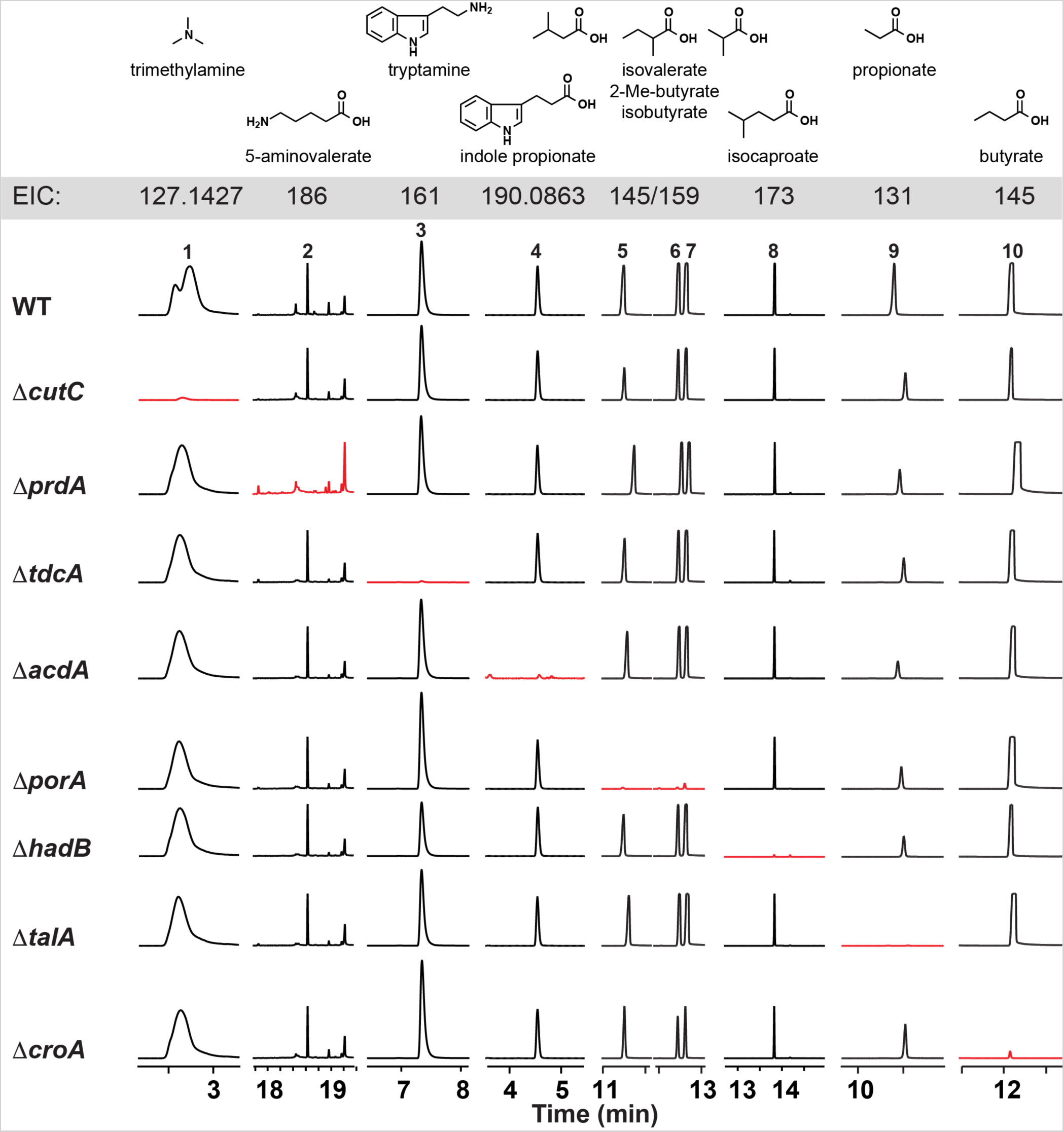
*Cs* mutants exhibit specific loss of metabolite production *in vitro*. Wild-type and mutant strains of *Cs* were cultured individually with the pathway substrates and metabolites were assayed by LC-MS and GC-MS. Extracted ion chromatogram windows corresponding to each of the pathway products are shown in order to compare the metabolic output of each strain; ion counts (y-axis) are on the same scale within a column, but have been scaled between columns. Full traces are displayed in **Figure S5**.Traces in red show the metabolite whose production is blocked by the mutation indicated at the beginning of each row. Each mutant is deficient in the production of the corresponding pathway product, but proficient in the production of all other pathway products.

### *In vivo* modulation of microbiome-derived small molecules using *Cs* mutants

Having validated our target pathways *in vitro*, we set out to determine whether we could use these mutants to “toggle” off the production of each pathway product in the context of host colonization. For this experiment, we used a subset of five mutants. The first four were *ΔcutC*, *ΔprdA*, *ΔcroA*, and *ΔhadB*; for the fifth, we took advantage of the markerless nature of our CRISPR-based genetic system to construct a *ΔporA/ΔhadB* double mutant, with the goal of eliminating the production of all four branched SCFAs (isobutyrate, 2-methylbutyrate, and isovalerate via *ΔporA*, and isocaproate via *ΔhadB*). We mono-colonized germ-free mice with WT *Cs* and the five mutants (**Figure 1**); after four weeks, we sacrificed the mice and measured the concentration of each molecule in serum, urine, cecal contents, and fecal pellets. We drew three observations from these data: 1) Each metabolite (or its host metabolic product) was substantially reduced in the corresponding mutant (**Figure 4**). 2) The WT/mutant differences were smaller for isocaproate and butyrate due to a combination of two factors: low production by WT *Cs* relative to the native level of each molecule (usually mM), and a background level of the metabolite in mutant-colonized mice, possibly due to a source of contaminating molecule in the chow. 3) The WT/mutant difference was especially large for the branched SCFAs isobutyrate, 2-methylbutyrate, and isovalerate, which form a pool of >2 mM in the cecal contents of WT *Cs*-associated mice that falls to near-baseline in the *ΔporA/ΔhadB*- associated animals. Overall, these data validate the utility and generality of using *Cs* mutants to deplete microbiome-derived molecules in the host, and they highlight that the presence of a pathway in an organism can lead to production levels that vary from native to orders of magnitude below. Thus, the choice of a producer organism that can support the biosynthesis of native metabolite levels depends on unknown factors beyond the mere presence of a corresponding pathway.

**Figure 4.**
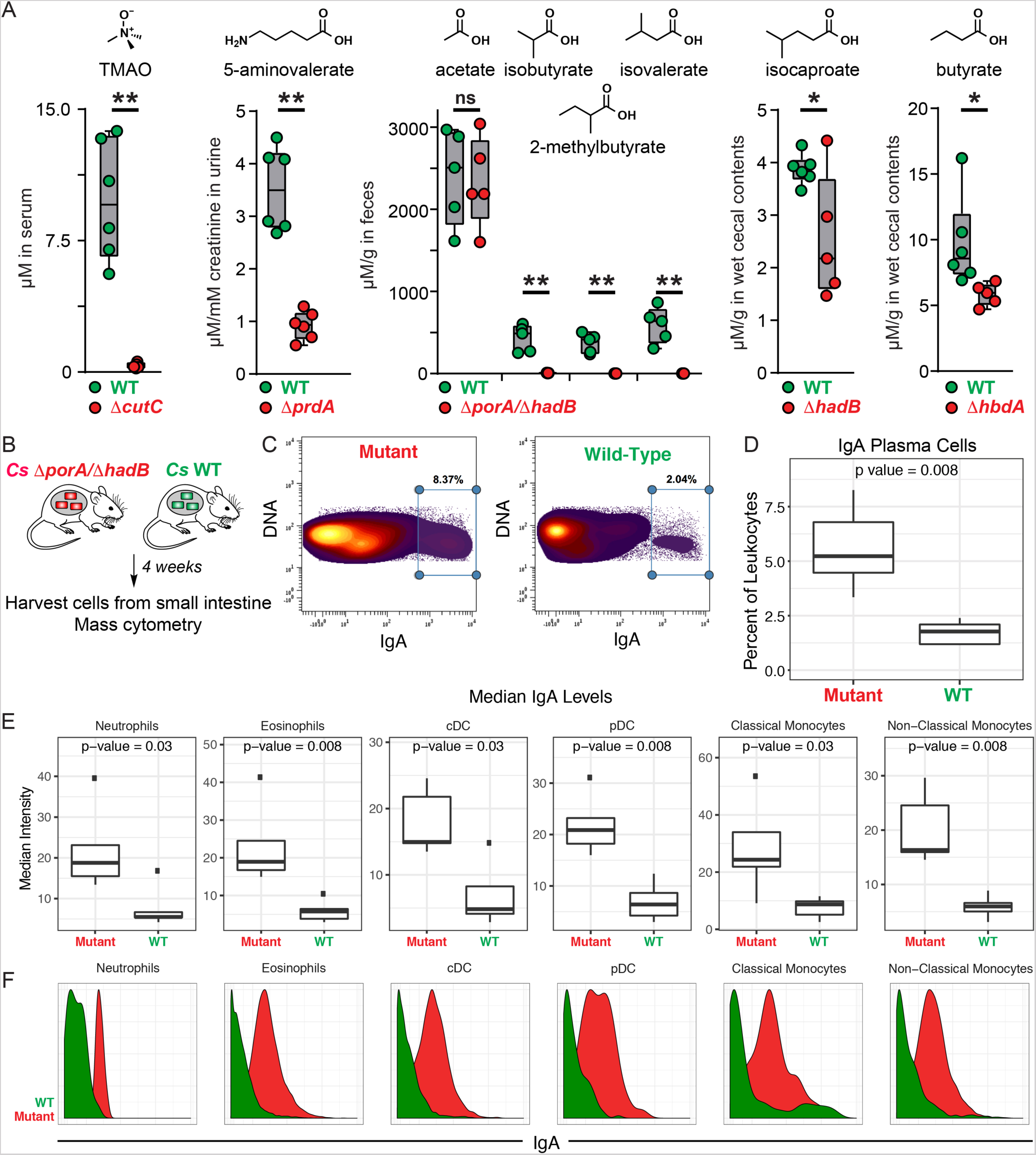
Genetic depletion of metabolites in vivo by colonization with wild-type vs mutant strains of *Cs*. (A). Germ-free mice were mono-colonized with wild-type *Cs* or the *ΔcutC, ΔprdA, ΔporA*/*ΔhadB,ΔhadB*, or *ΔhbdA* mutants. Metabolite levels were altered in the host by mutation of the corresponding pathway. (B) Schematic of germ-free mouse mono-colonization experiment. (C-F) Small intestinal lamina propria cells were analyzed by mass cytometry (n = 5 per group). (C) Frequency of IgA^hi^ cells of total live immune cells. (D) IgA plasma cells were quantified as a percent of live immune cells (CD45+) after excluding cells expressing common markers of other lineages (Ly6G, Siglec-F, B220, CD3, F4/80, CD64, CD11c, CD90, CD115, NK1.1, CD49b, Fc*ε*R1*α*). (E-F) Levels of surface-bound IgA were quantified on the indicated immune cell populations. Statistical analysis was performed using a Wilcoxon rank-sum test for all comparisons.

### Genetic control of microbiota metabolite production uncovers immune modulatory properties of BSCFAs

In light of our ability to ‘toggle’ off the *Cs*-derived small molecules *in vivo*, we returned to our original motivation for establishing the new genetic system: to study the role of microbiome-derived molecules in mediating microbe-host interactions. We chose the branched SCFAs for two reasons: (*i*) Like the conventional SCFAs acetate, propionate, and butyrate, they are highly abundant in the cecum and colon (concentrations in the mM range) (*26*); but unlike the SCFAs, which are known to modulate the host immune response via GPR41/43 (*1*), very little is known about the biology of the branched SCFAs. (*ii*) *Cs* produces them robustly; they constitute a pool of >2 mM (**Figure 4**). Hypothesizing that the branched SCFAs – like conventional SCFAs – might modulate the host immune response, we colonized germ-free mice with WT *Cs* and the *ΔporA*/*ΔhadB* mutant. After five weeks, we sacrificed the mice, isolated immune cells from the small intestine and mesenteric lymph nodes, and analyzed them by mass cytometry. WT- and *ΔporA*/*ΔhadB*-colonized mice were similar by broad metrics of immune function (e.g., total numbers of CD4+ and CD8+ T cells, B cells, and innate immune cells; percentages of effector cell subpopulations; cytokine levels). However, the *ΔporA*/*ΔhadB*-colonized mice had an increased number of immunoglobulin A (IgA)-producing plasma cells (**Figures 4D and S6**) and elevated levels of IgA bound to the surface of a variety of innate immune cells known to express IgA receptors: neutrophils, eosinophils, plasmacytoid and conventional dendritic cells, and classical and non-classical monocytes (**Figure 4E-F**). These results suggest a previously unrecognized role for the branched SCFAs in suppressing IgA production.

## Discussion

The hundreds of microbiome-derived molecules that accumulate in circulation represent one of the most concrete modes of communication between the host and its microbiota. Remarkably little progress has been made in systematically studying their effects on host biology, due in part to the absence of a method that enables the selective depletion of one member of this pool. One approach to addressing this challenge is to colonize germ-free mice with a wild-type gut bacterial species versus a metabolite-deficient mutant. Given the outsize role of *Clostridium* and related anaerobic Firmicutes in generating this pool of high-abundance metabolites, the lack of a reliable genetic system for commensal strains of *Clostridium* has been a key impediment to generalizing this approach. The system we introduce here is a first step toward that goal; it validates that genetics can be performed in a microbiome-derived *Clostridium* species rapidly, reliably, and without the need for a marker, and it demonstrates the utility of WT/mutant pairs for interrogating the host effects of microbiome-derived molecules. Given the versatility of Cas9, the key factors for generalizing this system to other *Clostridium* species are the ability to get DNA into a strain and the availability of replication origins for the plasmids that deliver the mutagenesis and repair elements. An alternative strategy would be to deliver Cas9 and the gRNA as a purified ribonucleoprotein, forgoing the need for plasmid-borne elements. Both strategies could expand the scope of Cas9-mediated mutagenesis to important but previously inaccessible Firmicutes such as butyrate producers from *Clostridium* clusters IV/XIVa (e.g., *Faecalibacterium prausnitzii*) (*27*), the bile acid metabolizers *C. scindens* and *C. hylemonae* (*28*), and the leanness-associated bacterium *Christensenella minuta* (*29*).

*Cs* is a prolific producer of amino acid metabolites, and our in vitro studies show that we can predictably block each pathway. However, one of the most striking observations is the difference in the concentration of each pathway product in the context of host colonization. 5-aminovalerate, indole propionate, isobutyrate, 2-methylbutyrate, isovalerate, propionate, and trimethylamine-*N*-oxide (TMAO) are produced robustly *in vivo* whereas isocaproate and butyrate are generated at 10-100-fold below the native concentration range. By highlighting that the mere presence of a pathway does not imply that it will function robustly in the host, our data raise questions about computational approaches that predict metabolic function based on gene or transcript levels (*30, 31*), and they suggest the importance of understanding pathway regulation and substrate availability under conditions of colonization by a complex native community.

The conventional SCFAs acetate, propionate, and butyrate modulate host immune function and induce regulatory T cells via a pair of GPCRs, GPR41/43. Though they are also predominantly microbiome-derived and present at high concentration (*26*), much less is known about the branched SCFAs isobutyrate, 2-methylbutyrate, and isovalerate. Isovalerate was recently shown to be a ligand for Olfr558, a GPCR in enterochromaffin cells that controls the secretion of serotonin (*32*). Our data uncover a novel role for the branched SCFA pathway in modulating the production of IgA, the predominant mucosally secreted antibody (*33*) whose mechanisms of microbiome modulation are an active area of study (*34–36*). In light of the fact that the conventional SCFAs have been shown to exert the opposite effect – induction of IgA-producing B cells (*37*) – it will be of interest to determine the molecular mechanism by which the branched SCFAs act, which could represent a novel point of control for a component of the adaptive immune response that is fundamentally important to host-microbe interactions at mucosal interfaces.

## ACKNOWLEDGMENTS

We are deeply indebted to members of the Fischbach Group for helpful suggestions and comments on the manuscript. This work was supported by an HHMI-Simons Faculty Scholar Award (M.A.F.); a Fellowship for Science and Engineering from the David and Lucile Packard Foundation (M.A.F.); an Investigators in the Pathogenesis of Infectious Disease award from the Burroughs Wellcome Foundation (M.A.F.); an award from BASF (M.A.F.); an award from the Leducq Foundation (M.A.F.); NIH grants DK110174 (M.A.F.), DK113598 (M.A.F.), DK101674 (M.A.F. and J.L.S.), OD023056 (M.H.S.), OD018040 (UCSF, to acquire the mass cytometer used in this study), DK110335 (D.D.), and DK085025 (J.L.S.); the Chan Zuckerberg Biohub (M.A.F. and J.L.S.); support from Stanford’s ChEM-H Institute (C.R.F.); and support from the Parker Institute for Cancer Immunotherapy (M.H.S.).

## SUPPLEMENTARY MATERIALS

## MATERIALS AND METHODS

### Assessing the metagenomic abundance of each pathway

We used MetaQuery (*1*) to assess the metagenomic abundance of each pathway. We started by identifying a key gene in each pathway – one that is predicted to encode an enzyme that catalyzes essential or committed step: *cutC* (trimethylamine), *prdA* (5-aminovalerate), *tdcA* (tryptamine), *fldB* (indolepropionate), *porA* (isovalerate, 2-methylbutryate, isobutyrate), *hadB* (isocaproate), *talA* (propionate), and *thlA* (butyrate). We used the following parameters in Metaquery: pairwise identity >50%, maximum e-value: 1e-5, query and target alignment coverage >70%, across 2000 publicly available human metagenomic stool samples. The results are shown in **Figure S2**.

### Assessing the metatranscriptomic abundance of each pathway

We built a local DNA sequence database that consists of 479 bacterial reference genomes obtained from the NIH Human Microbiome Project (HMP) website (https://hmpdacc.org/). We used the amino acid sequences of CutC, PrdA, TdcA, FldB, PorA, HadB, TalA, and ThlA as queries in a tblastn search of this database to identity homologs of each enzyme (maximum E-value: 1e-5). After removing hits with coverage <50% or pairwise identity <40%, we used the remaining sequences to construct a local database for metatranscriptomic analysis. We mapped metatranscriptomic reads from the stool of nine healthy human subjects (*2*) to this database using Bowtie 2 (local, medium sensitivity); representative mapping results are shown in **Figure S2**. We then set out to compare the expression of our target genes with its surrounding neighbors in the genome. For any gene to which reads mapped, we extracted a genomic region of ∼100 kb (∼50 kb 5’ of the gene and ∼50kb 3’) to construct a second database. The same metatranscriptomic dataset was mapped to this database using Bowtie 2 (local, medium sensitivity); representative results for two clusters, *prd* and *por*, are shown in **Figure S2**.

### Constructing *Cs* mutants using CRISPR/Cas9

#### Vector assembly

Primers and strains used in this study are listed in **Tables S1** and **S2**, respectively. The coding sequence of Cas9 was cloned from the vector pMJ825 (Addgene) using primers 83153_Cas9_(A10D)_XbaI_F and 83153_Cas9_XhoI_R. The purified PCR product and pMTL83153 were digested with XbaI/XhoI and ligated together using Instant Sticky-end Ligase (NEB), yielding pMTL83153_fdx_Cas9 (plasmid 1, P1) (**Figures 2** and **S3**).

We assembled DNA sequences encoding the gRNA locus (the gRNA plus adjacent elements) and the repair template into pMTL82254 as a backbone (specific details below). The repair template consists of two 700-1200 bp sequences flanking the 40-100 bp sequence targeted for excision. The design of the gRNA locus is shown below:

GTGCTACCAACACATCAA<u>GC**GGCGCC**</u>TTGACATGGGCTCACGAGAGCCTCTACTATAATATTGT TAGCTTGCCGTATACACAAGTTTTAGAGCTAGAAATAGCAAGTTAAAATAAGGCTAGTCCGTTATCAACTTGAAAAAGTGGCACCGAGTCGGTGCTTTTTTAGGAGAATAGAAAGAAGAAAATTCTTTCTAAAGGCTGAATTCTCTGTTTAATTTTGAGAGACCATTCTCTCAAAATTGAAACTTCTCAATAAAAATTGAGAAGTAGCTGACCATCACAAAATCGTAGATTTTGGATGTCTAGCTATGTTCTTTGAAAATTG CACAGTGAATAAGTAAAGCTAAAGGTATATAAAAATCCTTTGTAAGAATACAATTT**<u>GGCGCC</u>** <u>GC</u>AAAGTGACAGAGGAAAGC

The sequences highlighted in green are homologous to regions in pMTL82254. The sequence in blue is a synthetic promoter predicted by PePPER using the *Clostridium sporogenes* ATCC 15579 (*Cs*) genome as the calculation input (*3*). The red sequence is a small guide RNA (sgRNA) targeting the *Cs* chromosome. The sequence in yellow is for Cas9 binding, and the pink sequence is a terminator region obtained from the *Cs* 16s rRNA gene (CLOSPO_00916). The underlined sequences are NotI restriction sites that were not used in this study.

Using the *cutC* gene as an example, we used two primer pairs, 02864_TMA_F1+R2 and 02864_TMA_F3+R4 (**Table S1**), to synthesize the repair template, which consists of two regions (700-1200 bp each) flanking the sequence that is targeted for excision. The gRNA locus was synthesized commercially (gBlocks, IDT). The synthetic gRNA locus was fused to the repair template using fusion PCR with the primer set gRNA_F_NotI + gRNA_R_AscI. The purified PCR product (consisting of gRNA locus + repair template) was then cloned into a pMTL82254 backbone (*4*) that had been doubly digested by NotI/AscI using Instant Sticky-end Ligase (NEB), yielding pMTL82254_TMA_gRNA+rep temp (plasmid 2, P2) (**Figures 2** and **S3**). P1 and P2 (**Figure S3**) were introduced into two separate strains of *E. coli* S17 by electroporation.

#### Introducing vectors by conjugation into *Cs*

The process of constructing a single *Cs* mutant consists of two sequential conjugations from *E. coli* into *Cs.* For the first conjugation, a single colony of wild-type *Cs* was used to inoculate a 2 ml TYG broth culture in an anaerobic chamber at 37 °C under an atmosphere consisting of 10% CO_2_, 5% H_2_, 85% N_2_ *E. coli* S17 harboring P2 was grown in LB broth supplemented with erythromycin (250 µg/mL) at 30 °C with shaking at 225 rpm. After 17-24 hours, 1 mL *E. coli* S17 culture was centrifuged at 1000 × *g* for 1 min. The supernatant was discarded and the cell pellet was washed twice with 500 µL PBS buffer (pH = 7.2). The washed *E. coli* cell pellet was transferred into the anaerobic chamber, and 250 µL of *Cs* overnight culture was added and thoroughly mixed with *E. coli* cell pellet by pipetting. A 30 µL aliquot of the cell mixture was plated on a pre-reduced TYG agar plate as a liquid dot (a total of eight dots) for 24 h. Cell material in these dots was removed from the plate using a sterile inoculation loop and suspended in 250 µL pre-reduced PBS buffer (pH = 7.2). 100 µL of the cell suspension was plated on TYG agar + 10 µg/mL erythromycin + 250 µg/mL D-cycloserine. *Cs* colonies typically appeared after 36-48 h. Three colonies were picked and re-streaked on TYG agar + 10 µg/mL erythromycin + 250 µg/mL D-cycloserine to isolate single colonies. The transformation efficiency was typically high for this step, so one single colony was picked as the starting point for the second conjugation.

In the second conjugation, *E. coli* S17 harboring P1 was grown in LB broth supplemented with chloramphenicol (25 µg/mL) at 30 °C with shaking at 225 rpm. After washing the *E. coli* cell pellet as described in the previous paragraph, the washed *E. coli* cell pellet was transferred into anaerobic chamber and 250 µL of an overnight culture *Cs* (harboring vector P2) was added and thoroughly mixed with the *E. coli* cell pellet by pipetting. A 30 µL aliquot of the cell mixture was plated on a pre-reduced TYG agar plate as a liquid dot (a total of eight dots) for 72 h. Cell material in these dots was removed from the plate using a sterile inoculation loop and suspended in 250 µL pre-reduced PBS buffer (pH = 7.2). 100 µL of the cell suspension was plated on each of two TYG plates with 10 µg/mL erythromycin + 15 µg/mL thiamphenicol + 250 µg/mL D-cycloserine. *Cs* colonies typically appeared after 36-48 h. Sixteen colonies were picked and re-streaked on TYG agar + 10 µg/mL erythromycin + 15 µg/mL thiamphenicol + 250 µg/mL D-cycloserine to isolate single colonies. The isolated single colony was used to inoculate TYG broth supplemented with 10 µg/mL erythromycin + 15µg/mL thiamphenicol, and genomic DNA was isolated from the resulting cell material using Quick DNA fungal/bacterial kit (Zymo Research). We used diagnostic PCR and sequencing to identify mutants whose genomes harbor the desired deletions (see **Figure S4** for more details). For generating double-deletion mutants like Δ*porA*/Δ*hadB*, the first deletion was introduced using the CRISPR/Cas9 system. Following sequence verification, the mutant was plated on non-selective agar for multiple rounds to cure both plasmids, and then the process was repeated to introduce a second deletion.

### LC-MS/GC-MS analysis of *Cs* metabolites

A single colony of wild-type *Cs* or one of the mutant strains described herein was used to inoculate a 1 mL pre-reduced TYG broth culture [500 mL: 15 g tryptone, 10 g yeast extract, 0.5 g sodium thioglycolate, antibiotic concentration (if needed): thiamphenicol, 15 µg/ml; erythromycin, 10 µg/ml]. To pre-reduce, the TYG medium was left in the chamber with a loosened cap for at least 48 h before inoculation. The culture was incubated in an anaerobic chamber at 37 °C under an atmosphere consisting of 10% CO_2_, 5% H_2_, 85% N_2_.

#### 1) Quantification of the conversion of d_9_-choline to d_9_-trimethylamine (d_9_-TMA) by wild-type and mutant strains of *Cs*

<u>For in vitro bacterial cultures</u>: TYG broth was supplemented with 60 mM d_9_-choline. Following inoculation, the bacterial culture was incubated in the anaerobic chamber for 24 h. The overnight bacterial culture was centrifuged at 13000 × *g* for 10 min at room temperature. 100 µL of the cell-free supernatant was mixed with 10 µL of concentrated ammonia (7M in methanol) and 30 µL of ethyl bromoacetate (20 mg/mL in acetonitrile). The mixture was incubated at room temperature for ∼30 min and then quenched with equal volume of infusion solution (acetonitrile/water/formic acid, 50/50/0.025 (v/v/v)) (*5*).

1 µL of the quenched mixture was analyzed by LC-MS (Agilent 6530 QTOF) using the following conditions: The LC analysis was performed in positive mode using a Bio-Bond (Dikma Technologies) C4 column (5 μm, 4.6 mm × 50 mm), preceded by a C4 precolumn (3.5 μm, 2.0 mm × 20 mm). The mobile phase was 50/50 water/methanol (v/v) supplemented with 5 mM ammonium formate and 0.1% formic acid. The flow rate was 0.3 mL/min and run time was 6 min for each sample. The first 1.8 min of each analysis run was diverted to waste.

<u>For mouse samples</u>: To quantify the level of TMAO in serum samples, 20 µL serum was mixed with 80 µL of 10 µM d_9_-TMAO in MeOH, and 5 µL was analyzed using an Agilent 6470 LC-QQQ. The concentration of TMAO in the serum was determined by comparing its AUC (area under the curve) with that of d_9_-TMAO. We used the following chromatography conditions for the LC-QQQ: We used an Acquity UPLC BEH HILIC column (130 Å, 1.7 μm, 2.1 mm×100 mm, Waters Corp., Milford, MA, USA) with an Acquity UPLC BEH HILIC VanGuard pre-column (130 Å, 1.7 μm, 2.1 mm×5 mm). We used the following solvent system: A: H_2_O with 0.1% formic acid; B: Acetonitrile with 0.1% formic acid. 5 μL of each sample was injected, and the flow rate was 0.6 ml/min with a column temperature of 50 °C. The gradient for HPLC-MS analysis was: 0-0.8 min 5% A - 96% A, 0.8-1.9 min 96% A - 5% A. MRM was performed by filtering the precursor ions for m/z values of 76.1 to 58.1 (TMAO) and 85.1 to 68.1 (d_9_-TMAO), and the collision energies were 21 and 13 V, respectively (*6*). The y-axis of the graph depicting TMAO levels (at right in **Figure 4A**) was scaled to account for a previously identified difference in the intensity ratios of the two product ions between TMAO and d_9_-TMAO (*7*).

#### 2) Quantification of 5-aminovalerate production by wild-type and mutant strains of *Cs*

The sample preparation, derivatization, and chromatography conditions were adapted from a previously reported method (*8*). The bacterial culture was incubated in the anaerobic chamber for 48 hrs. Following incubation, a 100 µL aliquot of the culture was mixed with 80 µL of a propanol/pyridine solution (3:2, v/v), to which 10 µL propyl-chloroformate (PCF) was added. For urine samples, 20 µL of a urine sample was diluted 10-fold with ddH_2_O and mixed with 16 µL of a propanol/pyridine solution (3:2, v/v); 2 µL PCF was then added. The resulting derivatization reaction mixtures were sonicated at room temperature for 3 min and then extracted with an equal volume of hexanes. 1 µL of the organic layer was analyzed using a 7890B GC System (Agilent Technologies) and 5973 Network Mass Selective Detector (Agilent Technologies). We used the following chromatography conditions for GC-MS: Column: HP-5MS, 30 m, 0.25 mm, 0.25 µm; Injection volume: 1 µL; Injection Mode: splitless; Temperature Program: 40 °C for 0.1 min; 40-70 °C at 5 °C/min, hold at 70 °C for 3.5 min; 70-160 °C at 20 °C/min; 160 to 325 °C at 35 °C/min; equilibrate for 3 min.

#### 4) Detection of indole propionate production by wild-type and mutant strains of *Cs*

Following 24 hr incubation, the bacterial culture (1 ml total volume) was adjusted to pH ∼2 using 6M HCl and extracted 2x with an equal volume of ethyl acetate (EA). Solvent was removed from the combined EA fractions using a TurboVap. Indole propionate was validated by HRESIMS analysis [M - H]^-^ m/z found 188.0702, calcd for C_11_H_10_NO_2_ 188.0712]. The dried residue was resuspended in 100 µL 80%/20% DMSO/MeOH, and 5 µL of this solution was analyzed by LC-MS (Agilent 6530 QTOF) using the following conditions: Column: Agilent SB C-18, 1.8 µm, 3.0 × 100 mm; Solvent system: A: H_2_O with 0.1% formic acid; B: Acetonitrile with 0.1% formic acid. The gradient for LC-MS analysis was 0-5 min 100% A, 5-35 min 100% A - 0% A, 35-37 min 0% A, 37-39 min 0% A - 100% A, 39-41 min 100% A at a flow rate of 0.4 ml/min.

#### 5) Detection of isovalerate, 2-methylbutyrate, isobutyrate, isocaproate, propionate, and butyrate production by wild-type and mutant strains of *Cs*

Cultures were incubated for 48 h in the anaerobic chamber. Following incubation, a 50 µL aliquot of the culture was removed and acidified by the addition of 50 µL 6M HCl and 150 µL ddH_2_O. The resulting mixture was extracted with an equal volume of diethyl ether. For derivatization, 95 µl of diethyl ether extract was mixed with 5 µL N-tert-butyldimethylsilyl-N-methyltrifluoroacetamide (MTBSTFA) and incubated at room temperature for 48 h. Following derivatization, 1 µL of the organic layer was analyzed by GC-MS. Peaks were assigned by comparison with authentic standards and a standard curve was prepared for each chemical to quantify its concentration in biological samples. <u>For measuring propionate in wild-type</u> *<u>Cs</u>* <u>and the Δ</u>*<u>talA</u>* <u>mutant</u>: Both strains were grown in 5 mL TYG broth under anaerobic conditions for 48 h. Cultures were centrifuged at 5000 × g for 5 min, and the supernatant was discarded. The bacterial pellet was washed with pre-reduced PBS buffer (pH = 7.4, Gibco) twice, and then suspended in 1 mL PBS buffer. 100 µL L-threonine was added to the 900 µL cell suspension to reach a final concentration of 500 µM and incubated for 1 h, and a 50 µL aliquot was removed and acidified by the addition of 50 µL 6M HCl and 150 µL ddH_2_O. The resulting mixture was extracted with an equal volume of diethyl ether. For derivatization, 95 µl of diethyl ether extract was mixed with 5 µL MTBSTFA and incubated at room temperature for 48 h. Following derivatization, 1 µL of the organic layer was analyzed by GC-MS. <u>For samples from mice:</u> In brief, 500 µL of an extraction solution (10 µL 10 mM n-valeric acid in water as internal standard, 50 µL 6M HCl, 190 µL ddH_2_O, 250 µL diethyl ether) and six 6 mm ceramic beads were added to ∼100 mg wet cecal contents or fecal pellets. Samples were homogenized by vigorous shaking using a QIAGEN Tissue Lyser II at 25/s for 10 min. The resulting homogenates were subjected to centrifugation at 18000 × *g* for 10 min.

The organic layer was transferred to a new glass vial for derivatization using the following procedure: 95 µl of diethyl ether extract was mixed with 5 µL MTBSTFA and incubated at room temperature for 48 h. 1 µL of the derivatized samples were analyzed using a 7890B GC System (Agilent Technologies) and 5973 Network Mass Selective Detector (Agilent Technologies). We used the following chromatography conditions for GC-MS: Column: HP-5MS, 30 m, 0.25 mm, 0.25 µm; Injection Mode: splitless; Temperature Program: 50 °C for 2 min; 50-70 °C at 10 °C/min; 70-85 °C at 3 °C/min; 85 to 110 °C at 5 °C/min; 110 to 290 °C at 30 °C/min, equilibration for 3 min. 1 µL of each sample was injected and analyte concentrations were quantified by comparing their peak areas with those of authentic standards.

#### 6) Quantification of creatinine in urine samples

6 µL of a urine sample was mixed with 114 µLddH_2_O. The creatinine concentration of each diluted sample was measured using a Creatinine Assay Kit (ab204537).

### Colonizing germ-free mice with *Cs* strains

All experiments were performed using germ-free Swiss Webster mice (male, 6–10 weeks of age, n = 5 or 6 per group) originally obtained from Taconic Biosciences (Hudson, NY) and colonies were maintained in gnotobiotic isolators in accordance with A-PLAC, the Stanford IACUC. Germ-free mice were mono-colonized with wild-type or mutant strains of *Cs* by oral gavage of an overnight culture of WT or mutant *Cs* in TYG medium (200 µL; ∼1 × 10^7^ CFU).

For quantifying *Cs*-derived small molecules in vivo, mice (n = 5 or 6 per group) were maintained on standard chow (LabDiet 5K67). Urine and fecal samples were collected weekly and analyzed by GC-MS or LC-MS. After four weeks of colonization, mice were euthanized humanely by CO_2_ asphyxiation. Blood was collected by cardiac puncture and serum was prepared using microtainer serum separator tubes obtained from Becton Dickinson (Cat. # 365967). The urine, cecal contents, and feces were collected and snap-frozen in liquid nitrogen and stored at −80 °C until use.

For mass-cytometry analysis, two groups of germ-free mice (n = 5 per group) were mono-colonized by wild-type *Cs* or the Δ*porA*/Δ*hadB* double mutant by oral gavage. The mice were maintained on a high-protein chow (Teklad TD.90018; 40% protein) for four weeks. Fecal and urine samples were collected two weeks after colonization and analyte levels were measured using the procedures described above.

### Preparation of single-cell suspensions from the small intestine

Small intestines were harvested from animals and placed in RMPI medium on ice. Intestines were filleted open and washed 2x in PBS. Tissues were cut into 0.5 cm pieces and washed for 15 min in 5 ml HBSS + 0.015% DTT at 37 °C with steady rotation. Tissues were then washed for 30 min in 5 ml HBSS + 5% FCS + 25mM HEPES at 37 °C with steady rotation. Tissue pieces were rinsed in PBS thoroughly followed by complete RPMI medium and minced to ∼1 mm^3^ pieces with surgical scissors. Tissue pieces were digested in complete RPMI + 0.167 mg/ml Liberase (Roche) + 0.25 mg/ml DNAse I (Sigma-Aldrich) for 30 min at 37 °C. Digestion was quenched with complete RPMI, and cells were washed with PBS + 5 mM EDTA before labeling with 25 μM cisplatin viability die for mass cytometry analysis. Viability stain was quenched with PBS + 0.5% BSA + 25 mM EDTA. Cells were washed twice in PBS + 0.5% BSA + 0.02% NaN3 before fixation with 1.5% paraformaldehyde (Electron Microscopy Sciences) for 10 min at room temperature. Cells were washed twice in PBS + 0.5% BSA + 0.02% NaN3 and stored at −80 °C for subsequent mass cytometry analysis.

### Mass cytometry analysis

#### Antibody preparation

A summary of all mass cytometry antibodies, reporter isotopes and concentrations used for analysis can be found in **Table S3**. Primary conjugates of mass cytometry antibodies were prepared using the MaxPAR antibody conjugation kit (Fluidigm) according to the manufacturer’s recommended protocol. Following labeling, antibodies were diluted in Candor PBS Antibody Stabilization solution (Candor Bioscience GmbH, Wangen, Germany) supplemented with 0.02% NaN3 to between 0.1 and 0.3 mg/mL and stored long-term at 4 °C. Each antibody clone and lot was titrated to optimal staining concentrations using primary murine samples. One antibody cocktail was prepared for the staining of all samples for mass cytometry analysis.

#### Mass-Tag Cellular Barcoding

Mass-tag cellular barcoding was pre-formed as previously described (*9*). Briefly, 1*10^6^ cells from each animal were barcoded with distinct combinations of stable Pd isotopes in 0.02% saponin in PBS. All samples were barcoded together. Cells were washed two times in PBS with 0.5% BSA and 0.02% NaN3 and pooled into a single tube. After data collection, each condition was deconvoluted using a single-cell debarcoding algorithm (*9*).

#### Mass Cytometry Staining and Measurement

Cells were resuspended in PBS with 0.5% BSA and 0.02% NaN_3_ and unlabeled antibodies against CD16/32 were added at 20 μg/ml for 5 min at RT on a shaker to block Fc receptors. Surface marker antibodies were then added, yielding a 500 µL final reaction volume and stained at room temperature for 30 min at RT on a shaker. Following staining, cells were washed once with PBS with 0.5% BSA and 0.02% NaN_3_ then once with 1X Foxp3/Transcription factor permeabilization buffer (eBioscience). Cells were then stained with intracellular antibodies in a final volume of 500 μL permeabilization buffer for 30 min at RT on a shaker. Cells were washed twice in PBS with 0.5% BSA and 0.02% NaN_3_ and then stained with 1 mL of 1:4000 ^191/193^Ir DNA intercalator (Fluidigm) diluted in PBS with 1.6% PFA overnight. Cells were then washed once with PBS with 0.5% BSA and 0.02% NaN_3_ and then two times with double-deionized (dd)H_2_O. Care was taken to assure buffers preceding analysis were not contaminated with metals in the mass range above 100 Da. Mass cytometry samples were diluted in ddH_2_O containing bead standards (see below) to approximately 10^6^ cells per mL and then analyzed on a CyTOF^TM^ 2 mass cytometer (Fluidigm) equilibrated with ddH_2_O. We analyzed 2-6*10^5^ cells per animal, consistent with generally accepted practices in the field.

#### Bead Standard Data Normalization

Just before analysis, the stained and intercalated cell pellet was resuspended in ddH_2_O containing the bead standard at a concentration ranging between 1 and 2*10^4^ beads per ml as previously described (*10*). The bead standards were prepared immediately before analysis, and the mixture of beads and cells were filtered through a filter cap FACS tubes (BD Biosciences) before analysis. All mass cytometry files were normalized together using the mass cytometry data normalization algorithm (*10*), which uses the intensity values of a sliding window of these bead standards to correct for instrument fluctuations over time and between samples.

## SUPPLEMENTARY TEXT

### Developing a CRISPR/Cas9-based genetic system for *Cs*: Optimization and experimental design

We started by testing the CRISPR-Cas9 nickase system in *Cs*, as a similar system had been developed for *C. cellulolyticum*, a candidate for organic solvent production in industry (*11*). We assembled the Cas9 nickase, the small guide RNA (sgRNA), and a ∼2 kb repair template that flanks the targeted region in the *Cs* chromosome with pMTL83153 (*4*). We sequenced the *Cs* transconjugant and found no evidence of genome editing, leading us to try a genetic system with a fully functional Cas9 enzyme.

However, after replacing the Cas9 nickase with wild-type Cas9 in the assembled vector, we were unable to obtain any viable colonies that harbor the modified vector even after multiple trials. Since previous literature suggests that delayed or inducible Cas9 expression (*12, 13*) can improve genome editing efficiency, we put Cas9 under the control of either a *spoIIE* promoter or a lactose-inducible promoter. We still did not obtain any viable colonies with the resulting vectors.

In the meantime, we observed that we were unable to get viable colonies of *Cs* when the size of the conjugal vector exceeds ∼10 kb. To address this problem, we divided the components of the CRISPR-Cas9 system (Cas9 enzyme, gRNA, and repair template) between two separate vectors (**Figure S3**). However, we still failed to get viable colonies. By extending the incubation time of *Cs* with the *E. coli* conjugation donor to 72 h during conjugation, we were able to get one viable colony. Based on this observation, we developed an efficient protocol for mutating genes in *Cs* using CRISPR-Cas9 (See Materials and Methods for details).

## SUPPLEMENTARY FIGURES

**Figure S1.**
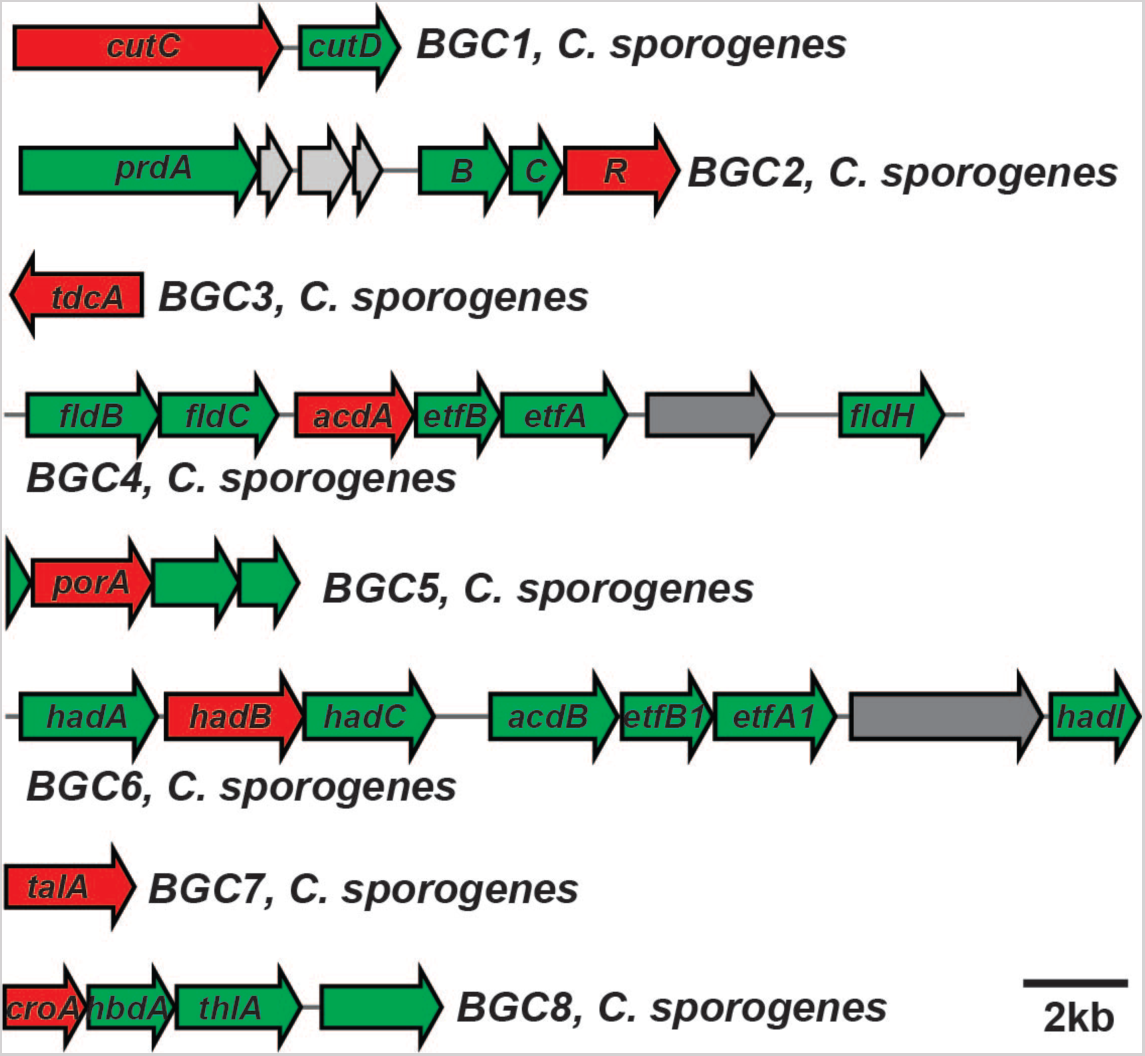
Gene clusters characterized in this study. A 40-100 bp fragment was deleted from each of the genes shown in red using the CRISPR/Cas9-based system.

**Figure S2.**
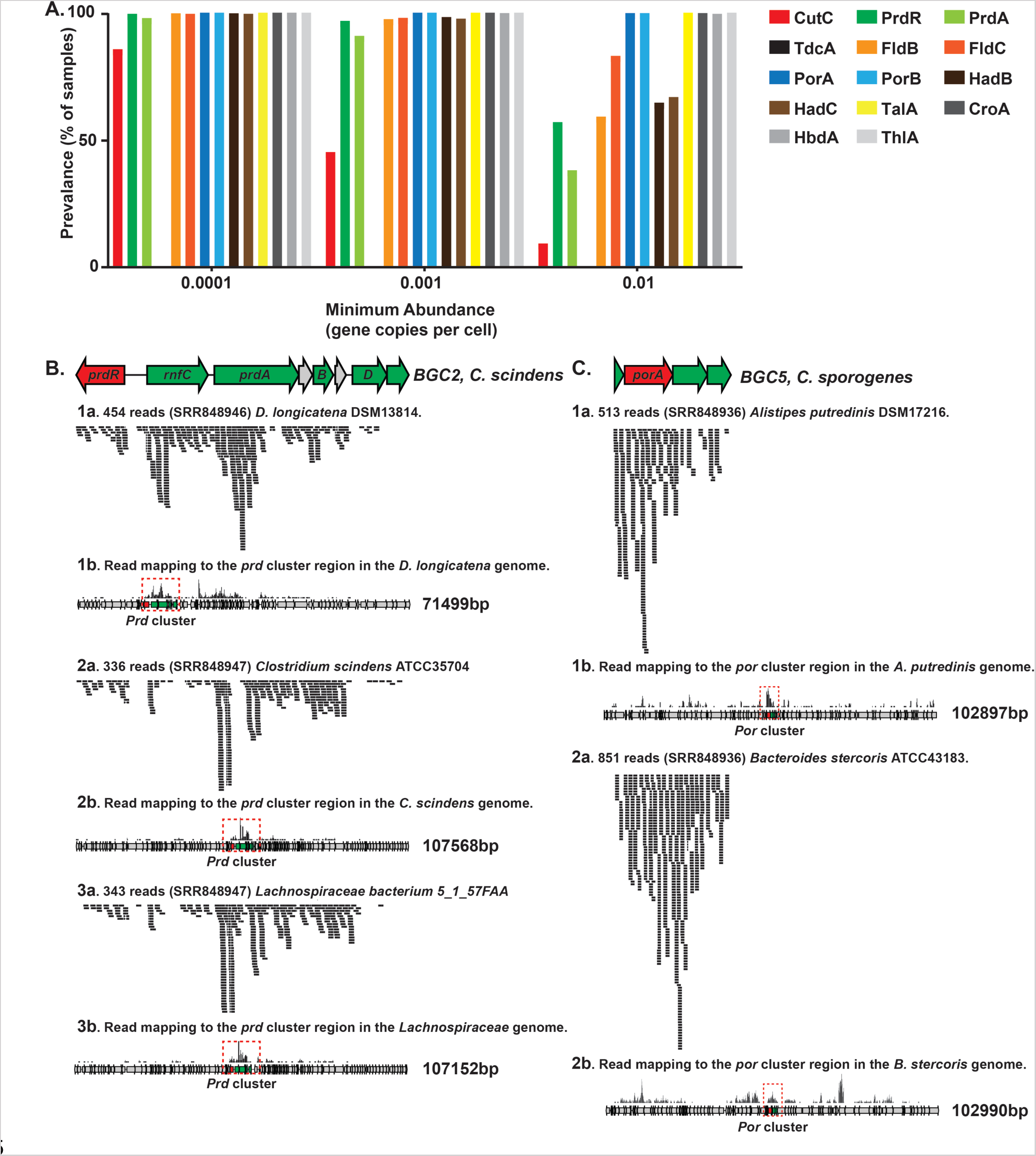
Metagenomic and metatranscriptomic analyses of the gene clusters and their homologs. (A) The key gene in each pathway – one that is predicted to encode an enzyme that catalyzes essential or committed step – was examined by MetaQuery to assess its metagenomic abundance across >2000 public available human stool metagenomes. (B-C) Representative results of metatranscriptomic analyses of MGCs and their homologs in this study. SRR8489XX indicates the accession number of each healthy human stool metatranscriptomic dataset examined in this study. For example, **Figure S2B, 1a** shows that 454 reads from the metatranscriptomic sample of human subject SRR848946 mapped to the *prd* cluster in *D. longicatena* DSM13814. This cluster is highly transcribed compared to the surrounding genes in the *D. longicatena* genome (**Figure S2B, 1b**).

**Figure S3.**
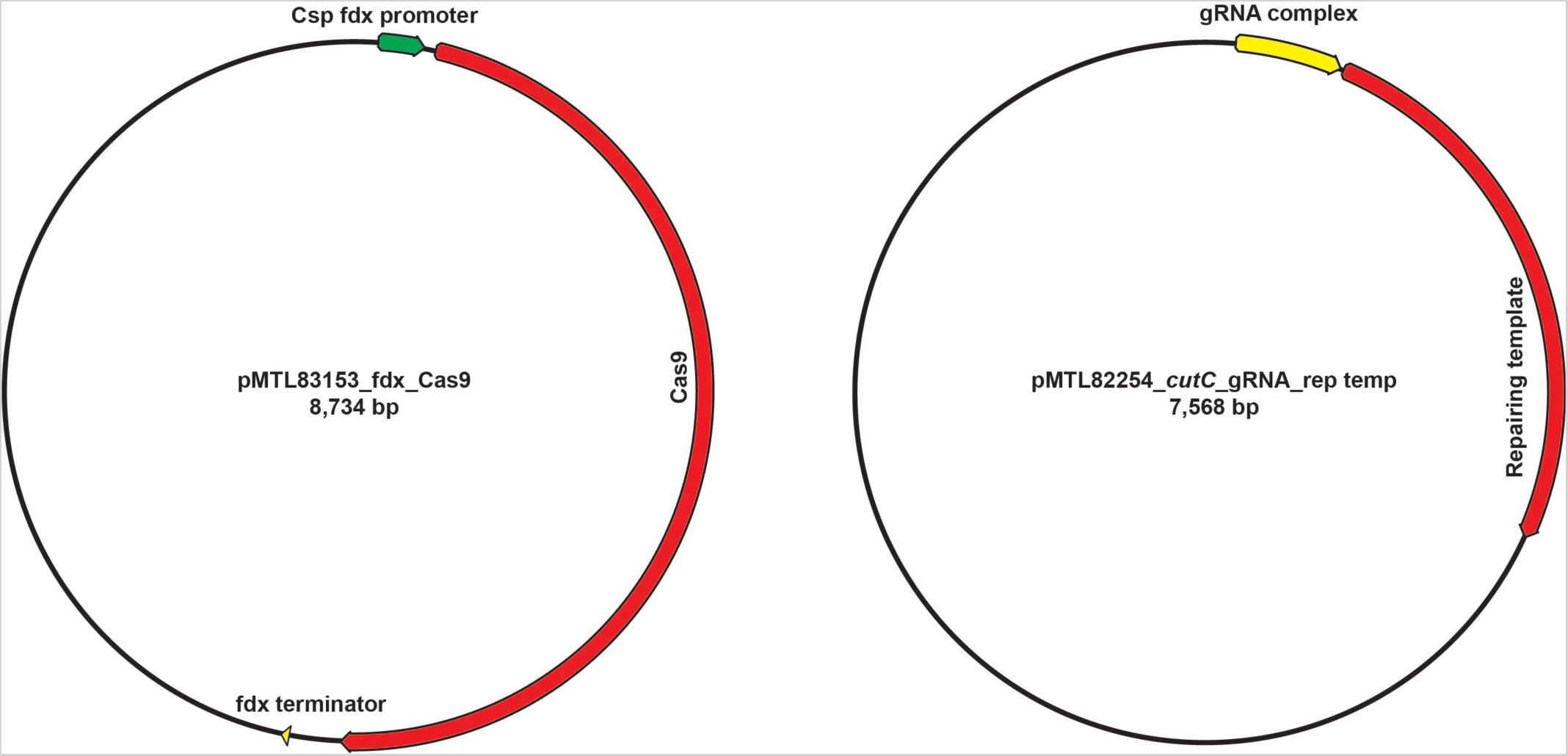
Schematics of two vectors designed in this study.

**Figure S4.**
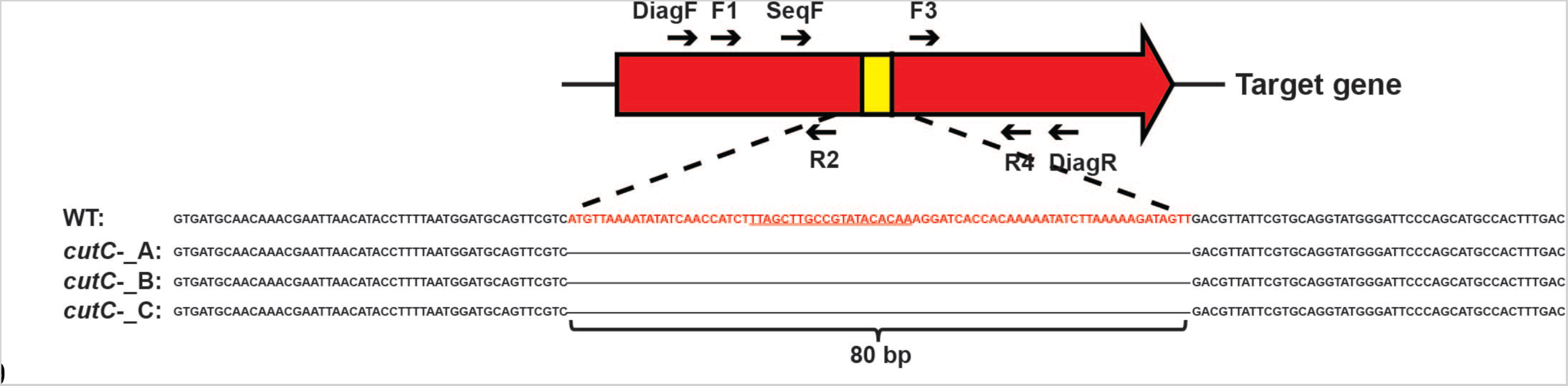
The scheme for diagnostic PCR used in this study and the sequencing result of three *cutC* mutants compared to wild-type *Cs*. The primer set DiagF+DiagR is used to amplify a sequence from the candidate mutants that carry both pMTL83153_fdx_Cas9 and pMTL82254_gRNA_rep Temp (**Figure S3**). The amplicons were then sequenced using the primer SeqF and compared to the sequence of WT *Cs* to determine whether the 80 bp sequence (in red) has been deleted in the mutant.

**Figure S5.**
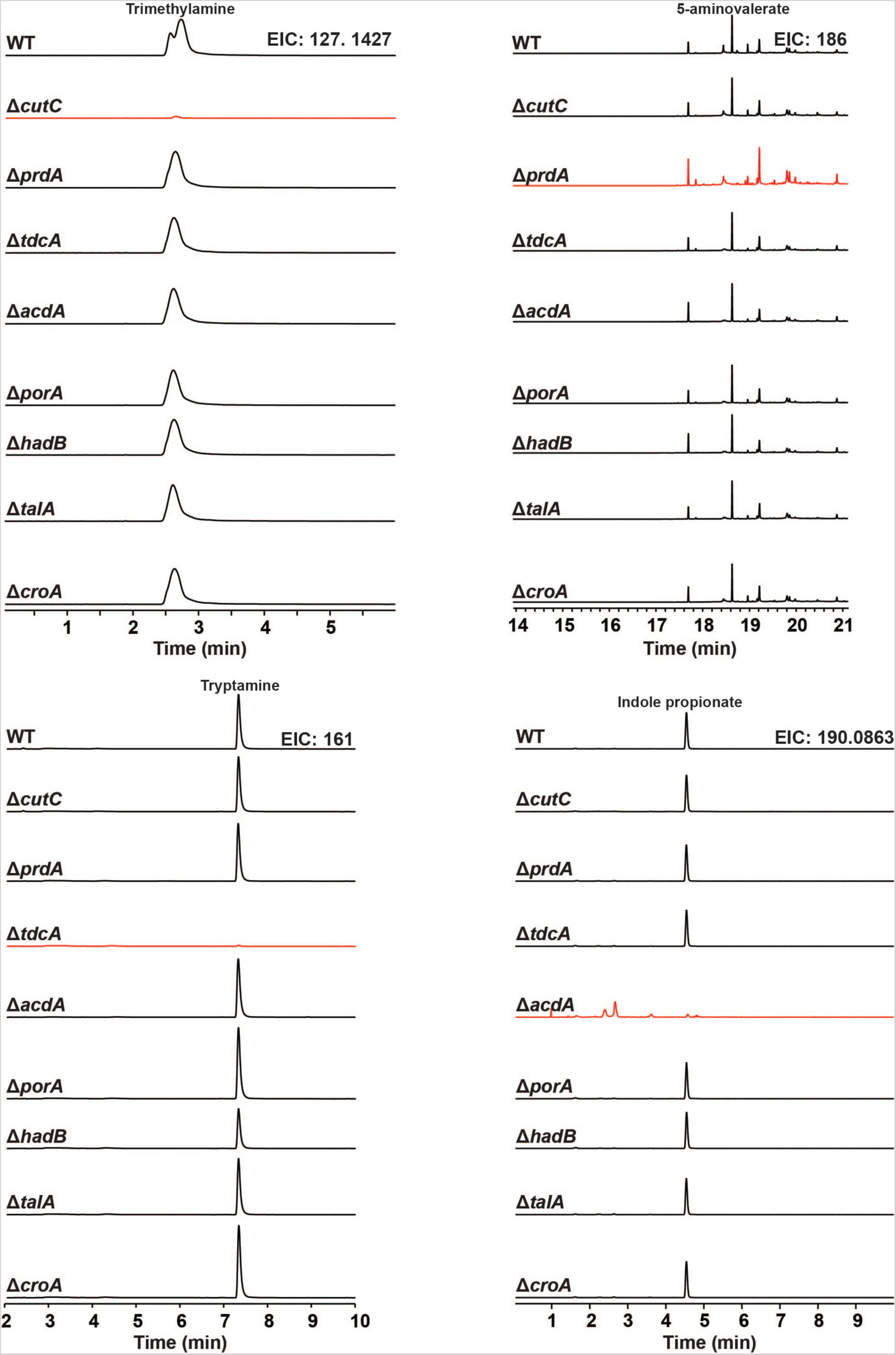
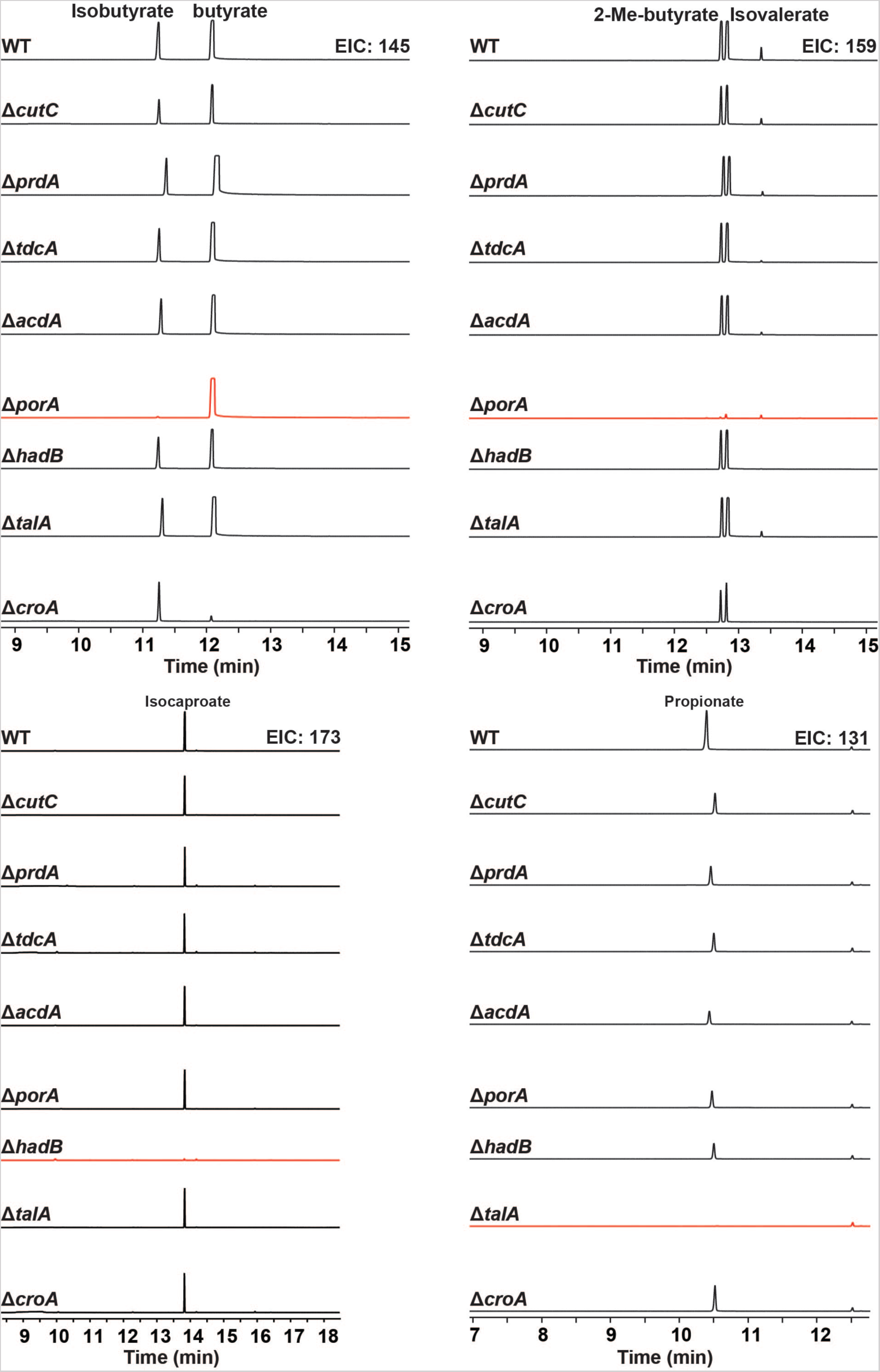
Complete MS chromatograms from wild-type and mutant strains of *Cs* cultured in vitro; a compressed version of these data is shown in **Figure 3**.

**Figure S6.**
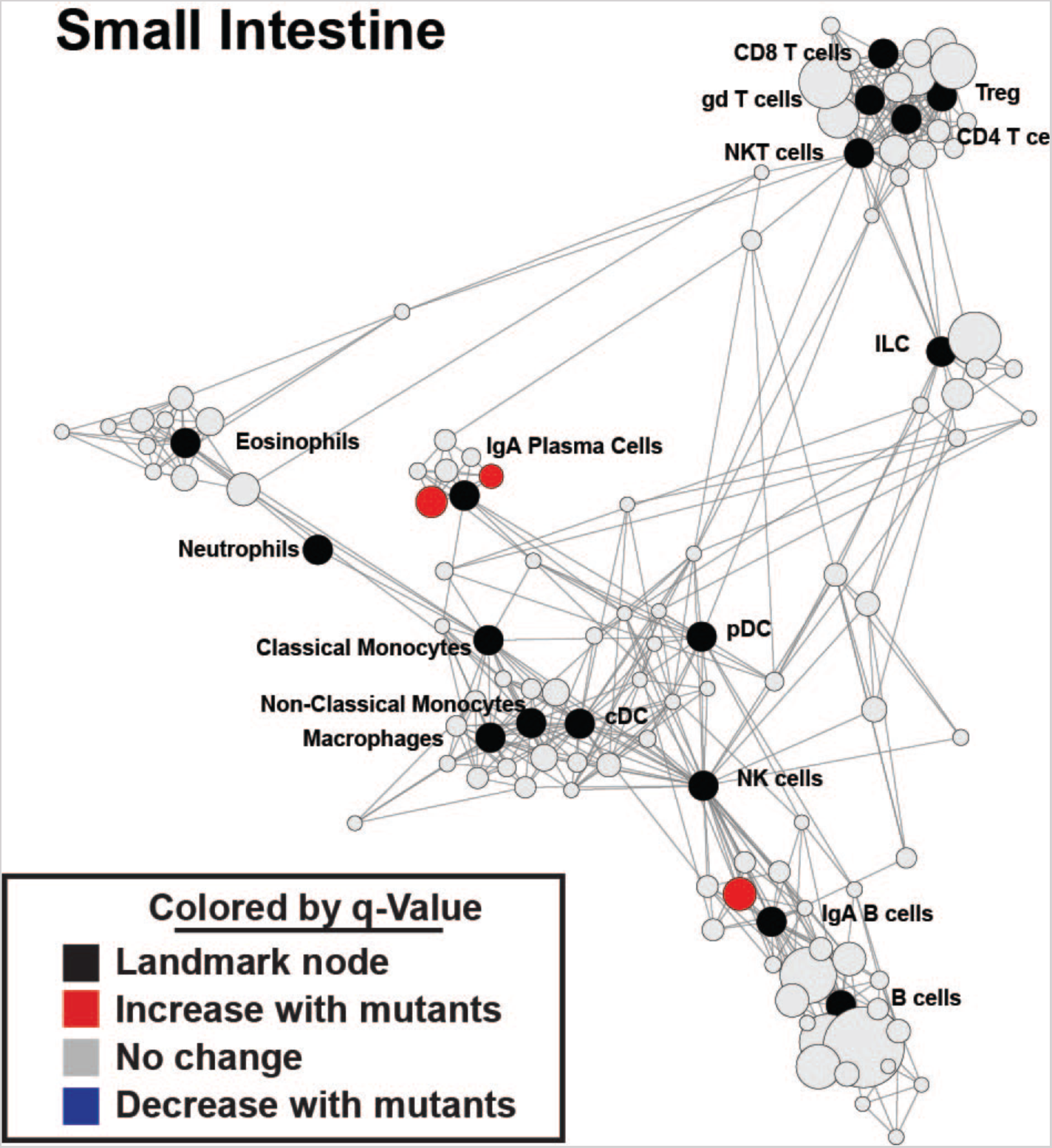
Fine detail scaffold maps of mass cytometry data. Scaffold maps of mass cytometry data from the small intestine and mesenteric lymph nodes of mice colonized with wild-type *Cs* or the ΔporA/ΔhadB mutant. Black nodes represent landmarks – canonical immune cell populations defined manually. Unsupervised cell clusters are positioned according to their nearest landmark node, with red nodes representing those immune cell populations that are upregulated in the mutant. The size of these nodes reflects the number of cells in that particular cluster. Edges in the graphs connect similar cells, with the length of each edge inversely proportional to that similarity. Cells that are most similar to one another are thereby connected by a short edge. The ΔporA/ΔhadB-colonized mice had an increased number of immunoglobulin A (IgA)-producing plasma cells as shown in the scaffold.

**Table S1.**
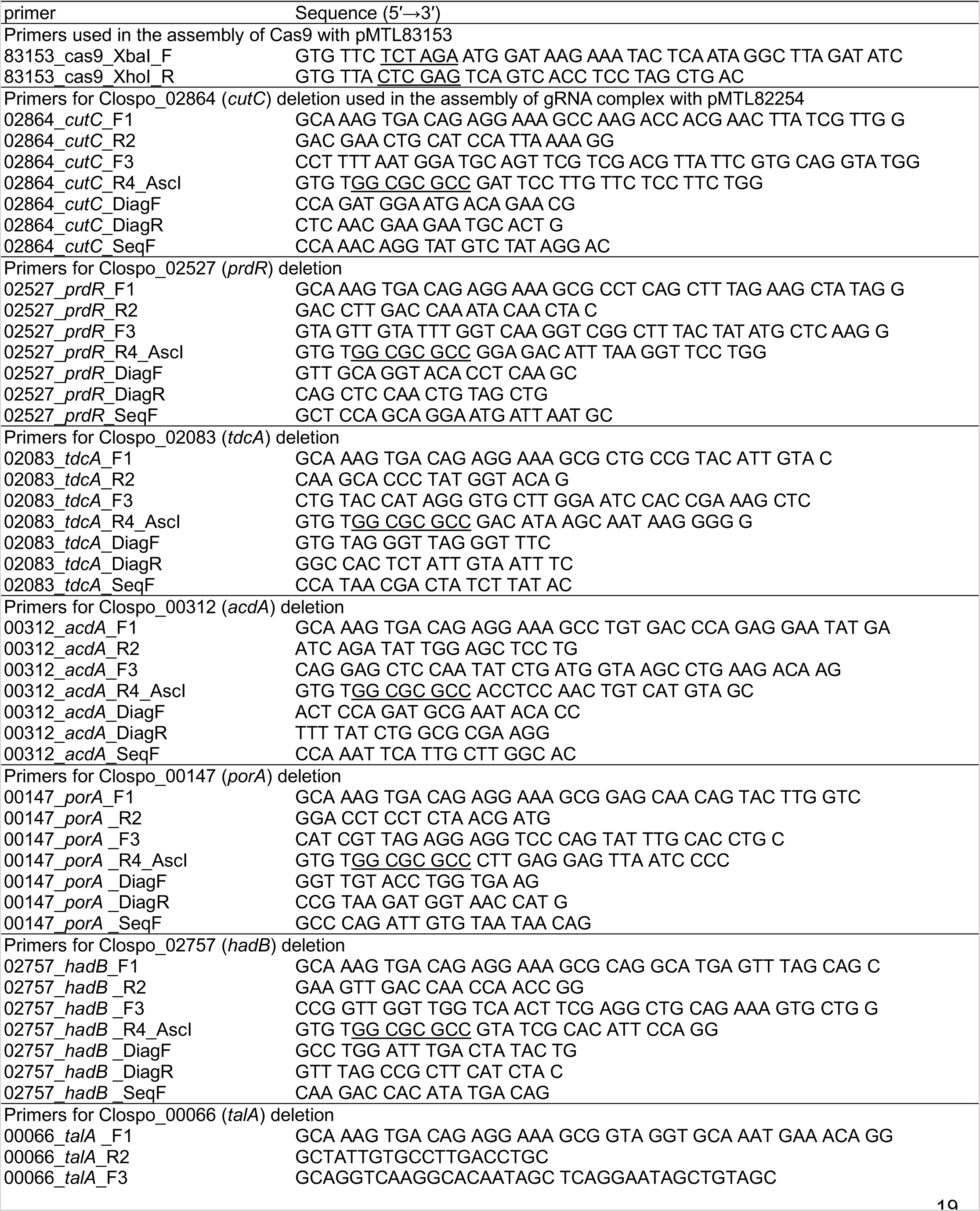
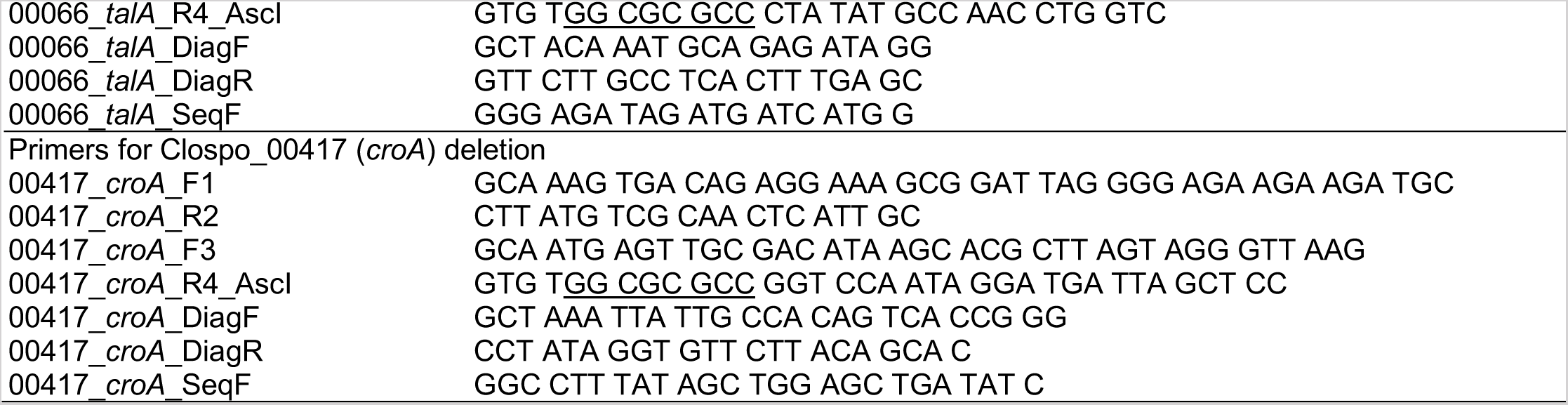
Primers used in this study.

**Table S2.**
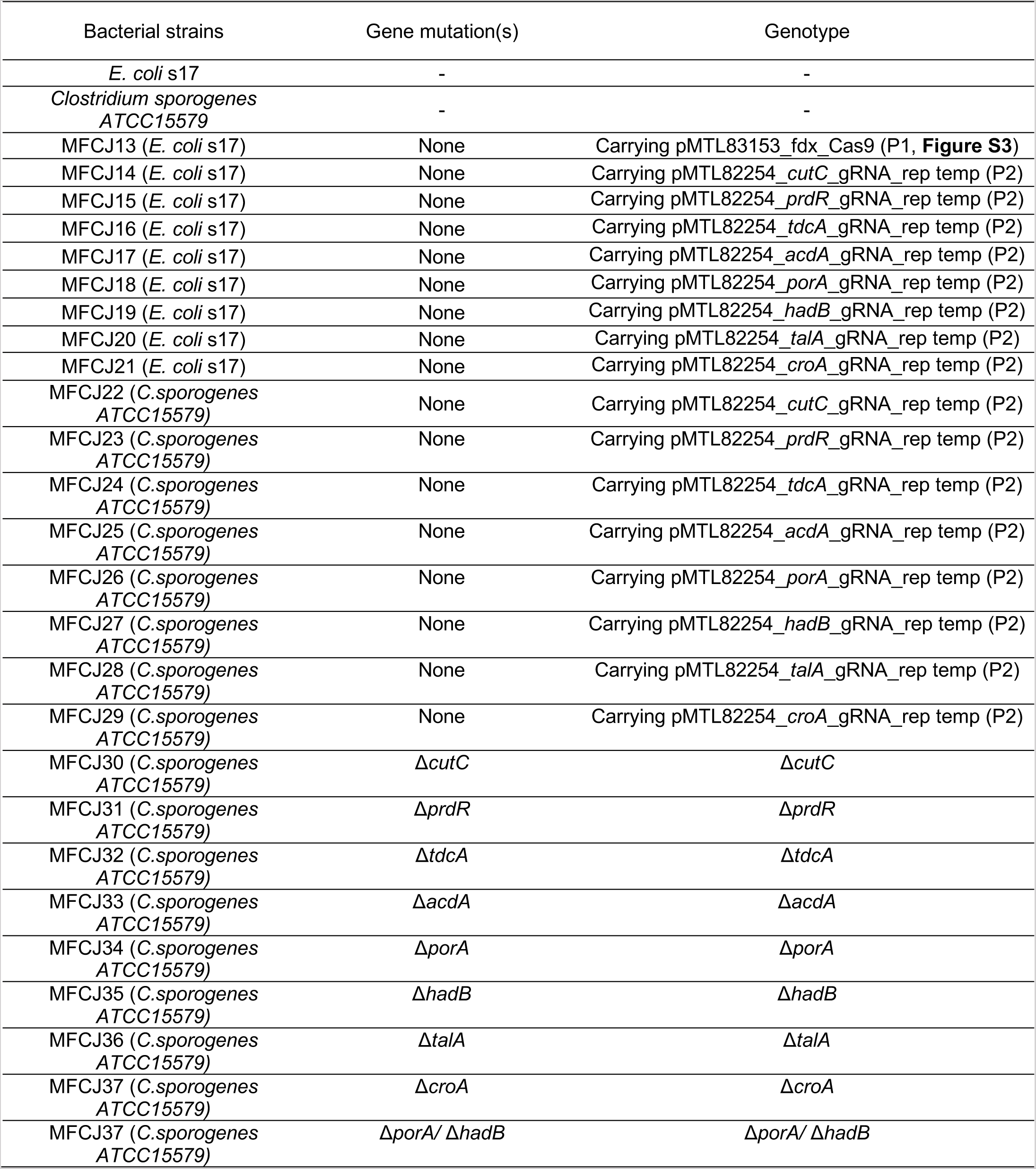
Bacterial strains used in this study.

**Table S3.**
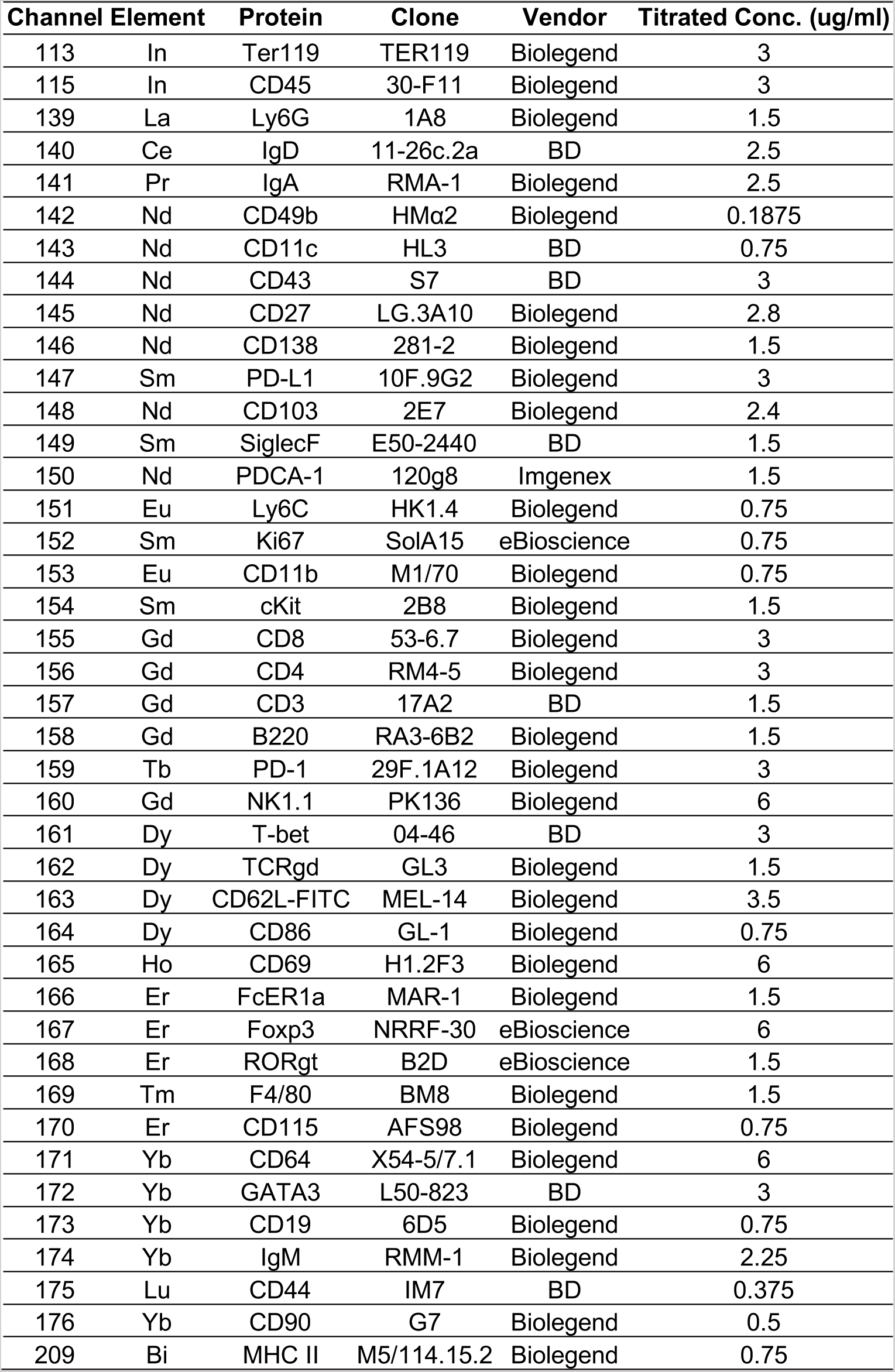
Antibody panel used for mass cytometry experiments.

**Table S4.**
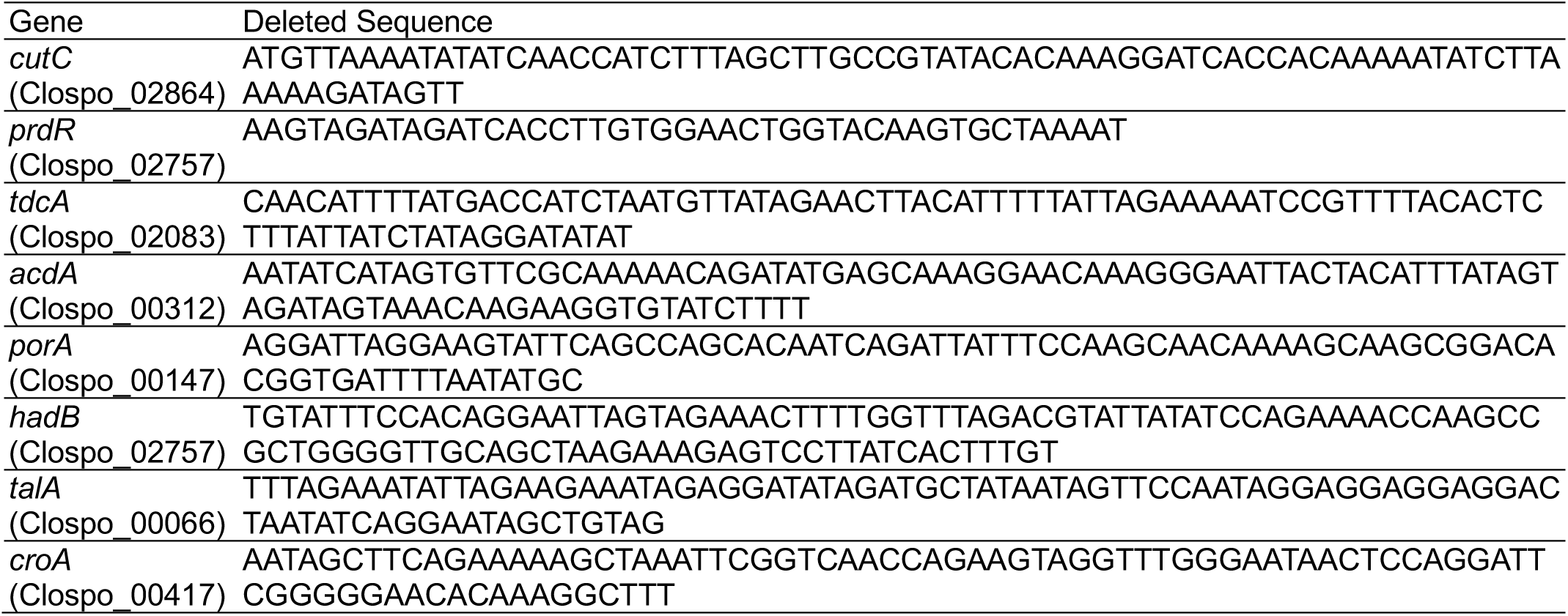
Gene regions deleted in this study.

## REFERENCES

1. M. G. Rooks, W. S. Garrett, Gut microbiota, metabolites and host immunity. Nat. Rev. Immunol. 16, 341–352 (2016).

2. P. M. Smith et al., The Microbial Metabolites, Short-Chain Fatty Acids, Regulate Colonic Treg Cell Homeostasis. Science (New York, N.Y. 341, 569–573 (2013).

3. Y. Furusawa et al., Commensal microbe-derived butyrate induces the differentiation of colonic regulatory T cells. Nature. 504, 446–450 (2013).

4. L. Zhao et al., Gut bacteria selectively promoted by dietary fibers alleviate type 2 diabetes. Science (New York, N.Y. 359, 1151–1156 (2018).

5. A. S. Devlin et al., Modulation of a Circulating Uremic Solute via Rational Genetic Manipulation of the Gut Microbiota. Cell Host & Microbe. 20, 709–715 (2016).

6. S. K. Mazmanian, H. L. Cui, A. O. Tzianabos, D. L. Kasper, An immunomodulatory molecule of symbiotic bacteria directs maturation of the host immune system. Cell. 122, 107–118 (2005).

7. K. A. Romano et al., Metabolic, Epigenetic, and Transgenerational Effects of Gut Bacterial Choline Consumption. Cell Host & Microbe. 22, 279–290.e7 (2017).

8. C. M. Thomas et al., Histamine derived from probiotic lactobacillus reuteri suppresses tnf via modulation of pka and erk signaling. PLoS ONE. 7 (2012), doi:10.1371/journal.pone.0031951.

9. S. A. Kuehne, J. T. Heap, C. M. Cooksley, S. T. Cartman, N. P. Minton, ClosTron-mediated engineering of Clostridium. Methods Mol. Biol. 765, 389–407 (2011).

10. D. Dodd et al., A gut bacterial pathway metabolizes aromatic amino acids into nine circulating metabolites. Nature. 551, 648–652 (2017).

11. S. R. Elsden, M. G. Hilton, J. M. Waller, The end products of the metabolism of aromatic amino acids by clostridia. Archives of microbiology. 107, 283–288 (1976).

12. S. R. Elsden, M. G. Hilton, Volatile acid production from threonin valine leucine and isoleucin by Clostridia. Archives of microbiology. 117, 165–172 (1978).

13. B. B. Williams et al., Discovery and Characterization of Gut Microbiota Decarboxylases that Can Produce the Neurotransmitter Tryptamine. Cell Host & Microbe. 16, 495–503 (2014).

14. A. Martínez-del Campo et al., Characterization and detection of a widely distributed gene cluster that predicts anaerobic choline utilization by human gut bacteria. MBio. 6, e00042–15 (2015).

15. U. C. Kabisch et al., Identification of D-proline reductase from Clostridium sticklandii as a selenoenzyme and indications for a catalytically active pyruvoyl group derived from a cysteine residue by cleavage of a proprotein. Journal of Biological Chemistry. 274, 8445–8454 (1999).

16. J. Kim, M. Hetzel, C. D. Boiangiu, W. Buckel, “Dehydration of (R)-2-hydroxyacyl-CoA to enoyl-CoA in the fermentation of α-amino acids by anaerobic bacteria” (2004), doi:10.1016/j.femsre.2004.03.001.

17. M. Vital, A. C. Howe, J. M. Tiedje, Revealing the Bacterial Synthesis Pathways by Analyzing (Meta) Genomic Data. MBio. 5, 1–11 (2014).

18. S. Nayfach, M. A. Fischbach, K. S. Pollard, MetaQuery: a web server for rapid annotation and quantitative analysis of specific genes in the human gut microbiome. Bioinformatics. 31, 3368–3370 (2015).

19. L. A. David et al., Diet rapidly and reproducibly alters the human gut microbiome. Nature. 505, 559–563 (2014).

20. J. T. Heap, O. J. Pennington, S. T. Cartman, N. P. Minton, A modular system for Clostridium shuttle plasmids. Journal of microbiological methods. 78, 79–85 (2009).

21. J. T. Heap, O. J. Pennington, S. T. Cartman, G. P. Carter, N. P. Minton, The ClosTron: A universal gene knock-out system for the genus Clostridium (2007), doi:10.1016/j.mimet.2007.05.021.

22. C. D. Denoya et al., A second branched-chain alpha-keto acid dehydrogenase gene cluster (bkdFGH) from Streptomyces avermitilis: its relationship to avermectin biosynthesis and the construction of a bkdF mutant suitable for the production of novel antiparasitic avermectins. Journal of bacteriology. 177, 3504–3511 (1995).

23. E. Chabrière et al., Crystal structures of the key anaerobic enzyme pyruvate ferredoxin oxidoreductase free and in complex with pyruvate. Nature Structural Biology. 6, 182–190 (1999).

24. J. Heider, X. Mai, M. W. Adams, Characterization of 2-ketoisovalerate ferredoxin oxidoreductase, a new and reversible coenzyme A-dependent enzyme involved in peptide fermentation by hyperthermophilic archaea. Journal of bacteriology. 178, 780–787 (1996).

25. K. Ma, A. Hutchins, S. J. Sung, M. W. Adams, Pyruvate ferredoxin oxidoreductase from the hyperthermophilic archaeon, Pyrococcus furiosus, functions as a CoA-dependent pyruvate decarboxylase. Proceedings of the National Academy of Sciences of the United States of America. 94, 9608–9613 (1997).

26. J. H. Cummings, E. W. Pomare, W. J. Branch, C. P. Naylor, G. T. MacFarlane, Short chain fatty acids in human large intestine, portal, hepatic and venous blood. Gut. 28, 1221–1227 (1987).

27. S. E. Pryde, S. H. Duncan, G. L. Hold, C. S. Stewart, H. J. F. Ã, The microbiology of butyrate formation in the human colon - 133.full.pdf. FEMS microbiology letters. 217, 133–139 (2002).

28. J. M. Ridlon, S. C. Harris, S. Bhowmik, D.-J. Kang, P. B. Hylemon, Consequences of bile salt biotransformations by intestinal bacteria. Gut Microbes. 7, 22–39 (2016).

29. J. K. Goodrich et al., Human Genetics Shape the Gut Microbiome. Cell. 159, 789–799 (2014).

30. M. G. I. Langille et al., Predictive functional profiling of microbial communities using 16S rRNA marker gene sequences. Nature biotechnology. 31, 814–821 (2013).

31. J. Kaminski et al., High-Specificity Targeted Functional Profiling in Microbial Communities with ShortBRED. PLOS Comput Biol. 11, e1004557 (2015).

32. N. W. Bellono et al., Enterochromaffin Cells Are Gut Chemosensors that Couple to Sensory Neural Pathways. Cell. 170, 185–198.e16 (2017).

33. A. J. MacPherson, K. D. McCoy, F. E. Johansen, P. Brandtzaeg, “The immune geography of IgA induction and function” (2008), pp. 11–22.

34. J. D. Planer et al., Development of the gut microbiota and mucosal IgA responses in twins and gnotobiotic mice. Nature. 534 (2016), doi:10.1038/nature17940.

35. A. L. Kau et al., Functional characterization of IgA-targeted bacterial taxa from malnourished Malawian children that produce diet-dependent enteropathy HHS Public Access. Sci Transl Med February. 25, 276–224 (2015).

36. J. J. Bunker et al., Natural polyreactive IgA antibodies coat the intestinal microbiota. Science (New York, N.Y. 358 (2017), doi:10.1126/science.aan6619.

37. M. Kim, Y. Qie, J. Park, C. H. Kim Correspondence, Gut Microbial Metabolites Fuel Host Antibody Responses. Cell Host & Microbe. 20, 202–214 (2016).

38. A. de Jong, H. Pietersma, M. Cordes, O. P. Kuipers, J. Kok, PePPER: a webserver for prediction of prokaryote promoter elements and regulons. BMC Genomics. 13 (2012), doi:10.1186/1471-2164-13-299.

## REFERENCES

1. S. Nayfach, M. A. Fischbach, K. S. Pollard, MetaQuery: a web server for rapid annotation and quantitative analysis of specific genes in the human gut microbiome. Bioinformatics. 31, 3368–3370 (2015).

2. L. A. David et al., Diet rapidly and reproducibly alters the human gut microbiome. Nature. 505, 559–563 (2014).

3. A. de Jong, H. Pietersma, M. Cordes, O. P. Kuipers, J. Kok, PePPER: a webserver for prediction of prokaryote promoter elements and regulons. BMC Genomics. 13 (2012), doi:10.1186/1471-2164-13-299.

4. J. T. Heap, O. J. Pennington, S. T. Cartman, N. P. Minton, A modular system for Clostridium shuttle plasmids. Journal of microbiological methods. 78, 79–85 (2009).

5. S. Craciun, E. P. Balskus, Microbial conversion of choline to trimethylamine requires a glycyl radical enzyme. Proceedings of the National Academy of Sciences of the United States of America. 109, 21307–21312 (2012).

6. L. M. Heaney, D. J. L. Jones, R. J. Mbasu, L. L. Ng, T. Suzuki, High mass accuracy assay for trimethylamine N-oxide using stable-isotope dilution with liquid chromatography coupled to orthogonal acceleration time of flight mass spectrometry with multiple reaction monitoring. Analytical and Bioanalytical Chemistry. 408, 797–804 (2016).

7. Z. Wang et al., Measurement of trimethylamine-N-oxide by stable isotope dilution liquid chromatography tandem mass spectrometry. Anal. Biochem. 455, 35–40 (2014).

8. X. Zheng et al., A targeted metabolomic protocol for short-chain fatty acids and branched-chain amino acids. arXiv. 9, 818–827 (2013).

9. E. R. Zunder et al., Palladium-based mass tag cell barcoding with a doublet-filtering scheme and single-cell deconvolution algorithm. Nat Protoc. 10, 316–333 (2015).

10. R. Finck et al., Normalization of mass cytometry data with bead standards. Cytometry A. 83, 483–494 (2013).

11. T. Xu et al., Efficient genome editing in clostridium cellulolyticum via CRISPR-Cas9 nickase. Applied and environmental microbiology. 81, 4423–4431 (2015).

12. Y. Wang et al., Markerless chromosomal gene deletion in Clostridium beijerinckii using CRISPR/Cas9 system. Journal of biotechnology. 200, 1–5 (2015).

13. Y. Wang et al., Bacterial Genome Editing with CRISPR-Cas9: Deletion, Integration, Single Nucleotide Modification, and Desirable “Clean” Mutant Selection in Clostridium beijerinckiias an Example. ACS Synth. Biol. 5, 721–732 (2016).

